# CetZ1-dependent polar assembly of the archaeal motility machinery

**DOI:** 10.1101/2024.05.02.592137

**Authors:** Hannah J. Brown, Md Imtiazul Islam, Juanfang Ruan, Matthew A. B. Baker, Solenne Ithurbide, Iain G. Duggin

## Abstract

The archaeal tubulin-like cytoskeletal protein CetZ1 is required for rod-cell morphogenesis during the development of motility in *Haloferax volcanii*. This is expected to improve swimming speed and directionality. Here, we found that deletion of *cetZ1* or expression of a GTPase-defective mutant caused a substantial defect in the assembly of the motility machinery, including the archaellum base marker protein ArlD1, the chemotaxis sensory array adapter CheW1, and signal transducer CheY. Furthermore, overexpression of *cetZ1* reduced the assembly and polar placement of the motility machinery without detectably affecting the rod shape of motile cells. In contrast, deletion of the conserved paralog *cetZ2* caused no defects in swimming or rod shape, although expression of the *cetZ2* GTPase-defective mutant reduced motility whereas *cetZ2* overexpression caused mild hyper-motility; these effects were dependent on the presence of *cetZ1*. A functional CetZ1-mTq2 fusion strongly localized at the poles of mature motile cells, where it partially co-localized with the motility machinery markers. These results suggest that CetZ1 has another role in the organisation or structure of the cell poles that promotes the assembly of the motility machinery. The multiple roles and locations of CetZ1 during motile cell development are reminiscent of the multiple functions of eukaryotic cytoskeletal proteins.

## Introduction

Cytoskeletal proteins are utilised as platforms or scaffolds for large protein complexes across the tree of life. Tubulin superfamily proteins are particularly widespread, including tubulins in Eukarya, FtsZ in Bacteria and Archaea, and the Archaea-specific CetZ family (Aylett and Duggin, 2017; Erickson, 1995; Erickson, 2007; Nogales et al., 1998; Vaughan et al., 2004). In eukaryotes, tubulin forms microtubules, which act as platforms in several biological contexts, including for the anchoring and partitioning of chromosomes during cell division (Barton and Goldstein, 1996), intracellular transport of cargo (Appert-Rolland et al., 2015; Hirokawa et al., 2009), and assembly of protein complexes (An et al., 2010) and surface appendages (Hawkins et al., 2010). In bacteria, FtsZ forms a ring at mid-cell which is an essential platform required for the assembly of the proteins at the division site, consisting of more than thirty proteins (Lutkenhaus et al., 2012). In comparison to tubulin and FtsZ, CetZs are understudied in terms of their basic biological functions and capacity to act as platforms or scaffolds for assembly of other protein complexes.

In the motile and pleomorphic model halophilic archaeon, *Haloferax volcanii*, CetZs have been shown to control cell shape changes and have also been implicated in swimming motility (de Silva et al., 2024; Duggin et al., 2015). *H. volcanii* has six CetZ paralogues, with CetZ1 and CetZ2 being the most highly conserved. CetZ1 is required for rod-development and similar morphological changes that occur in several conditions, including trace metal starvation, early log-phase growth, and when cells become motile in soft agar (de Silva et al., 2021; Duggin et al., 2015). Deletion of *cetZ1* and mutation of its GTPase active site (*cetZ1*.E218A), which is critical for the CetZ1 polymerisation cycle, both result in inhibition of rod-development and motility (de Silva et al., 2024; Duggin et al., 2015). Deletion of *cetZ2* has not been found to result in a cell shape or motility phenotype during standard growth conditions (Duggin et al., 2015), although expression of a *cetZ2* point mutant with an inactivated GTPase (*cetZ2*.E212A) prevents rod formation and causes a motility defect in a dominant inhibitory manner (Duggin et al., 2015). In addition, a survey of the presence of CetZs, and motility and rod shape phenotypes across Haloarchaea found that the presence of both CetZ1 and CetZ2 is highly correlated with motility and rod-shape (Brown and Duggin, 2023).

How might CetZ1 bring about its effects on motility? Fundamentally, the rod-shape of motile cells is expected to provide a faster, more streamlined hydrodynamic shape and improve the directionality of movement during swimming (Duggin et al., 2015; Young, 2006). Additionally, rod-shaped cells have a natural polarity which is a likely cue for the assembly and placement of structures at the poles, such as the motility machinery, through the influence of spatial regulators such as MinD family proteins. Previous work showed that *H. volcanii* MinD4 oscillates along the long axis of *H. volcanii* cells and dynamically localises at the cell poles, which influences the correct positioning of the motility proteins ArlD1 and CheW1 at or near the poles (Nußbaum et al., 2020). The paralog MinD2 was also recently implicated in the polar localisation ArlD1, CheW1, and CheY (Patro et al., 2024), and both MinD2 and MinD4 influence CetZ1 localization (Brown and Duggin, 2024). Furthermore, in plate-shaped cells, which do not have the well-defined polarity of rods, the MinD4 wave-like patterns tend to breakdown or move in circles instead of oscillating (Nußbaum et al., 2020). In plate-shaped stationary phase cells, the archaellum filament and chemosensory arrays are not localized, although the archaellum base protein ArlD1 is observed in foci (Li et al., 2019; Nußbaum et al., 2020). These studies suggest a significant association between rod shape and the assembly of the polar motility complexes.

CetZ1 might also have other functions that in motility that are not directly linked to its function in generating the rod shape (Brown and Duggin, 2023). Initial subcellular localization studies with a CetZ1-GFP fusion (Duggin et al., 2015) indicated that it assembles as dynamic filaments or sheet-like structures at the cell envelope as motile rod cells take shape, and then forms a cap or patch-like structures at the cell poles later in rod development. In mature motile rods sampled from motility agar, the polar CetZ1 structures appeared especially bright and stable at the poles. Polar cap-like structures have been previously characterized in archaea as large cytoplasmic structures which are situated adjacent to the inner membrane in motile euryarchaea; they have been identified in *Pyrococcus furiosus* (Daum et al., 2017), *Thermococcus kodakarensis* (Briegel et al., 2017), and *Halobacterium salinarum* (Kupper et al., 1994). The polar caps were found to act as polar organising centres associated with archaella bundles. In *P. furiosus* and *T. kodakarensis*, polar caps were also associated with chemosensory arrays. The proteins that likely comprise polar caps are not yet known. While a polar cap has not yet been identified in *H. volcanii*, the potential involvement of CetZ1 in recruiting, assembling or comprising a polar cap or the motility machinery raises another potential way that CetZ1 could promote motility.

Like in bacteria, archaeal chemotaxis arrays are responsible for sensing extracellular signals and activating a signal cascade that results in a change in rotational direction of the archaellum (Li et al., 2019; Li et al., 2020; Quax et al., 2018), which is the whip-like structure equivalent to the bacterial flagellum (reviewed in (Albers and Jarrell, 2015; Albers and Jarrell, 2018; Beeby et al., 2020; Jarrell and Albers, 2012; Jarrell et al., 2021)). Chemotaxis proteins in Haloarchaea are like those of the bacterial system, with methyl-accepting chemotaxis proteins (MCPs), CheW, and CheA organised as a chemosensory array at the cell envelope. When extracellular signals bind to the MCPs they induce a signal cascade that results in autophosphorylation of CheA, which in turn phosphorylates CheY (Li et al., 2020). Phosphorylated CheY shuttles between chemosensory arrays and the archaellum base (comprised of ArlD and ArlCE) where it binds and phosphorylates CheF (Li et al., 2019; Quax et al., 2018). Together, ArlD and ArlCE are responsible for signal transduction to the archaellum motor proteins (ArlH, ArlI, ArlJ), and they are highly dependent on these motor proteins for localization at the cell poles (Li et al., 2020). ArlD and ArlCE are thought to directly bind the archaellum motor (Briegel et al., 2017; Daum et al., 2017) but are also presumed to bind a hypothesized cytoplasmic cone- or cap-like structure at the cell poles (Li et al., 2020). In this study, we utilized a range of genetic and microscopy techniques to investigate the potential for CetZ1 and CetZ2 to contribute to motility via influencing cell shape and the assembly and positioning of polar archaellum and chemotaxis complexes

## Results

### CetZ1 and CetZ2 differentially influence motility in H. volcanii

CetZ1 has previously been implicated in cell shape and motility (Duggin et al., 2015), but it is not yet clear whether CetZs contribute to motility solely through their control of cell shape or in other ways too. To investigate this, we quantified the effects of deletion and overexpression of *cetZ1* and its paralog *cetZ2* on motility and motile-cell shape. We used soft-agar motility assays, where cells swim out from a central inoculation site at a characteristic rate, forming a halo (Fig. 1a). To assess motile cell shape, we withdrew cells from the leading edge of the halo for microscopy, and measured cell circularity as an assessment of relative cell elongation of the motile cells (Fig. 2a, 2b), as well as a comprehensive set of other rod cell shape characteristics, described below (Fig. 2c-i).

**Figure 1.**
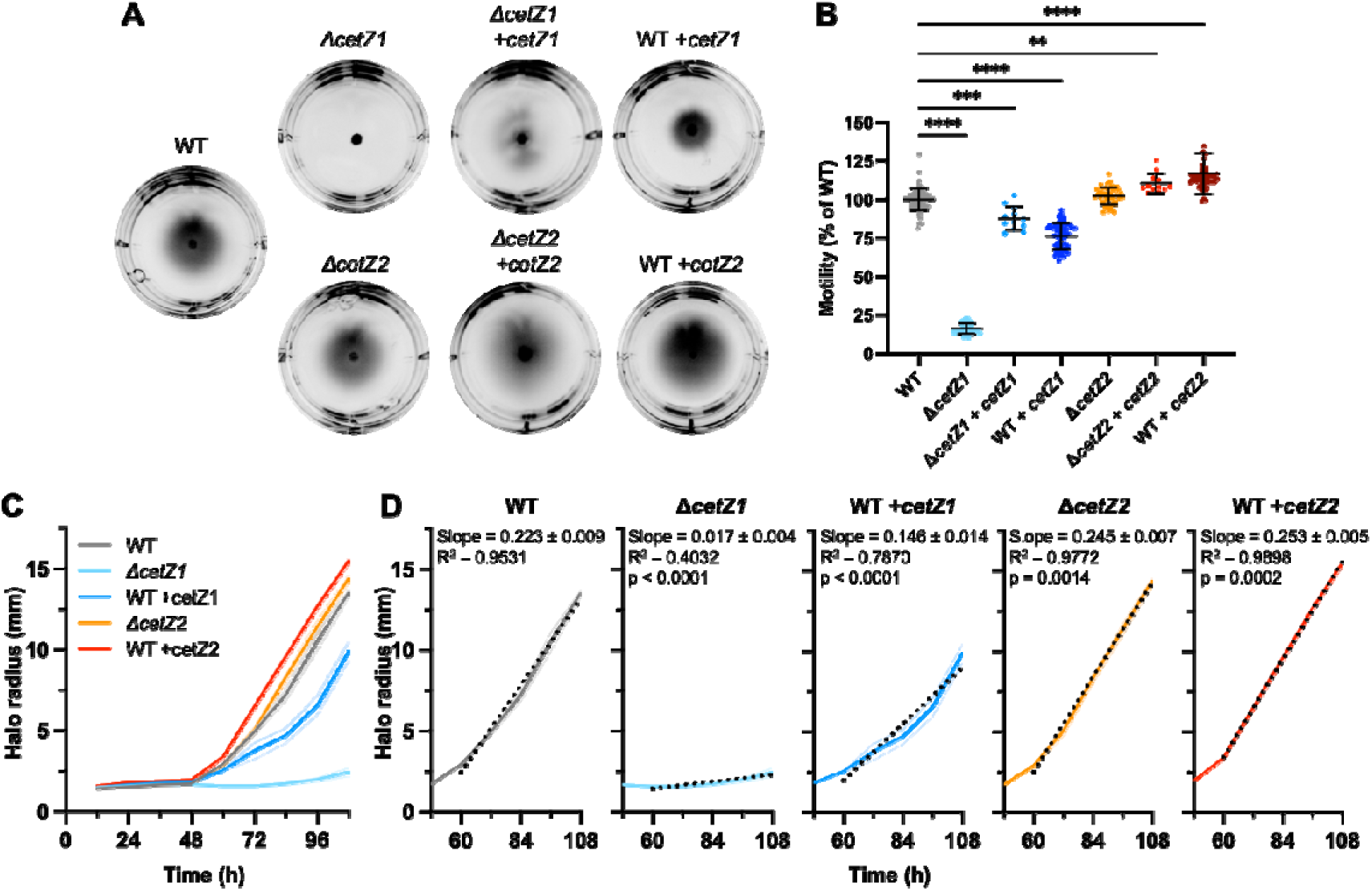
Comparison of motility amongst *cetZ1* and *cetZ2* mutants. **A)** H26 wildtype (WT), Δ*cetZ1* and Δ*cetZ2* backgrounds with pTA962 vector only, and H26 wildtype, Δ*cetZ1*, and Δ*cetZ2* with *cetZ1* or *cetZ2* expressed from pTA962 as indicated (+*cetZ1/2*) were inoculated onto Hv-Cab soft-agar with 1 mM L-Tryptophan as inducer. **B)** Their motility halo diameters were quantified after four days and normalized to the mean of H26. Individual points represent independent replicates. One-way ANOVA was used as a statistical test (****p<0.0001, ***p<0.0002, **p<0.002). **C)** Halo radius was quantified in 12-hour intervals. The bold line indicates the mean of six replicate halos, and shading represents the standard deviation. **D)** A linear regression (black dotted line) was fitted for halo radius measurements (data as in **C**) between 60 and 108 hours. The slope of the fitted line represents the mean halo expansion rate in mm/h and is noted in each graph with standard error of the slope and R^2^. A t-test was used as a statistical test to compare slopes of *cetZ1* and *cetZ2* mutant strains with H26 wildtype, the p-values of which are also noted.

**Figure 2.**
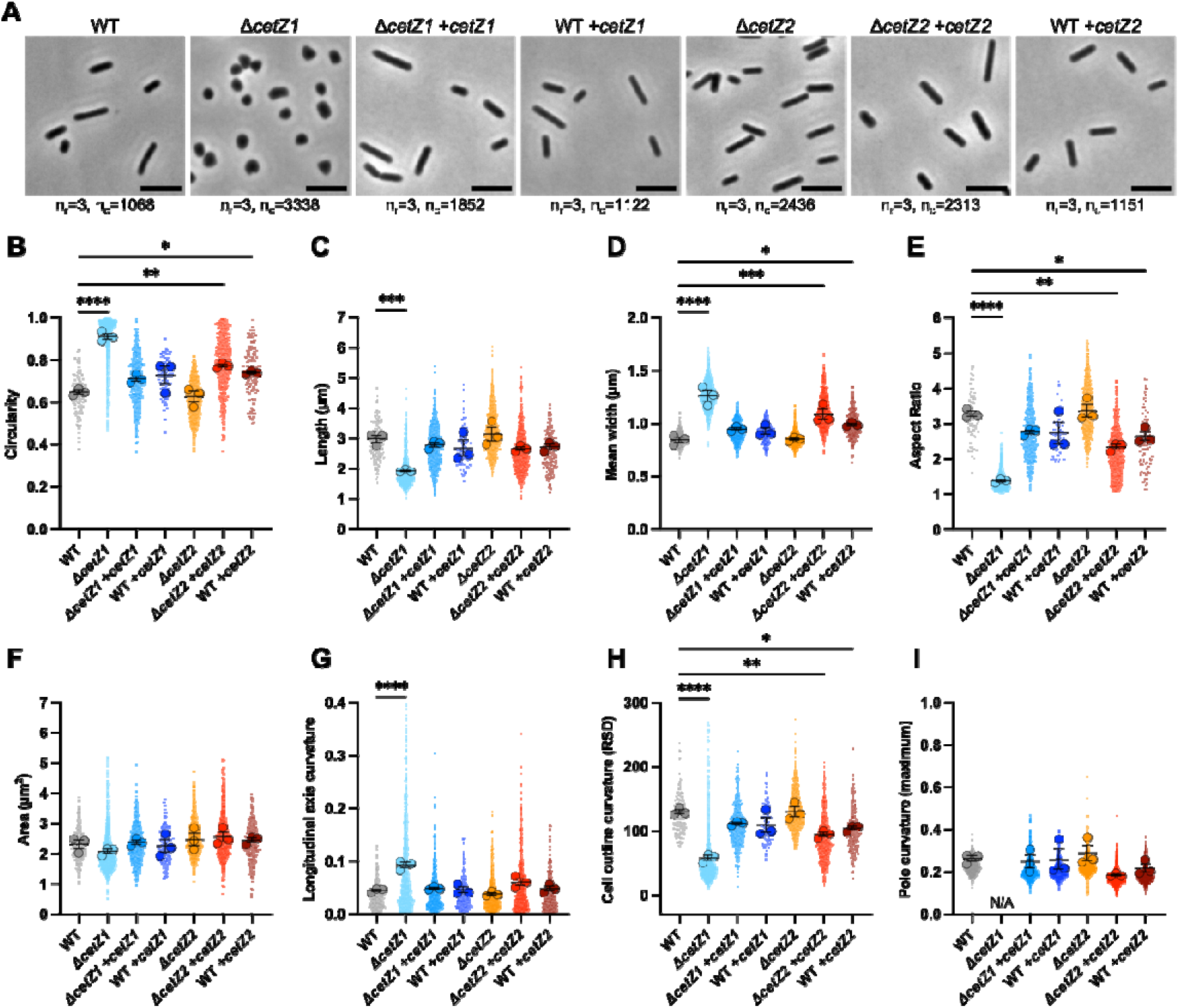
Comparison of motile cell shapes amongst *cetZ1* and *cetZ2* mutants. **A)** H26 wildtype (WT), Δ*cetZ1* and Δ*cetZ2* backgrounds with pTA962 vector only, and H26 wildtype, Δ*cetZ1*, and Δ*cetZ2* with *cetZ1* or *cetZ2* expressed from pTA962 as indicated (+*cetZ1/2*) were withdrawn from the edge of motility halos and imaged using phase-contrast microscopy. Scale bar: 5 μm. The number of cells (n_c_) pooled from all independent replicates (n_r_) for each strain are reported below each image and apply to all panels. Images were used to quantify various cell shape parameters, including **B)** circularity, **C)** cell length, **D)** mean width, **E)** aspect ratio, **F)** cell area, **G)** curvature of the cell longitudinal axis, **H)** relative standard deviation of the curvature of the cell outline, and **I)** maximum curvature of cell poles. For **B-H**, small data points indicate individual cells and for **I**, small data points indicate individual poles. For **B-I**, large data points indicate the mean of each biological replicate, and error bars indicate mean and standard error of biological replicate means. One-way ANOVA was used as a statistical test (****p<0.0001, ***p<0.0002, **p<0.002, *0.02), and only significant and/or relevant comparisons are shown.

As expected, deletion of *cetZ1* caused a severe motility defect (∼12% of wildtype motility, Fig. 1b) and prevented rod-formation in cells extracted from the leading edge of motility halos (Fig. 2a, 2b), consistent with previous work on a different *cetZ1* deletion and genetic background, H98 (Duggin et al., 2015). Deletion of *cetZ2* was previously reported to have no effects on motility or rod shape (Duggin et al., 2015), which was also confirmed here in motile cells (Fig. 1b, 2a, 2b). By expressing *cetZ1* from a plasmid in the Δ*cetZ1* background (ID622), we observed strong complementation of motility and motile-cell shape (Fig. 1b, 2b, S1, 2).

Interestingly, overexpression of *cetZ1* in the wildtype background using the tryptophan-inducible promoter (p.*tna*) on a plasmid based on pTA962 (Allers et al., 2010) caused a dose-dependent reduction in motility compared to the wildtype control (Fig. 1b, S3). We also discovered that overexpression of c*etZ2*, whether in wildtype or Δ*cetZ2* backgrounds, resulted in a mild increase in motility, approximately 115% of wildtype (Fig. 1b, S1a).

We further investigated the altered motility phenotypes of *cetZ1/2* deletion and overexpression strains by measuring the halo radii over time at 12-hour intervals. This showed that the onset of motility halo expansion occurred consistently after a delay of ∼48 hours after inoculation (Fig. 1c, d). During this 48 h delay period, we observed only localized colony growth at the inoculation site, inferring that the development of motile cells occurs in this colony during the delay period. The differences in halo size observed after 3 days (Fig. 1a, 1b) are therefore primarily due to differences in their radial expansion (net swimming) rates, and not potential differences in the onset delay period and motile cell differentiation. The wild-type halos expanded at a mean rate of 223 ± 0.009 µm/hour, which represents the overall (net) speed of *H. volcanii* swimming under these conditions. The *cetZ* mutant strains all showed statistically significant differences in radial expansion rate compared to the wildtype (Fig. 1d; compare slopes), which were consistent with the differences observed in the single time-point assays (Fig. 1b).

In the case of *cetZ1* deletion and *cetZ2* overexpression, the differences in overall swimming speed were accompanied by measurable changes in cell elongation, measured by cell circularity (Fig. 2a, b). However, we observed no measurable differences in cell circularity of the *cetZ1* overexpression strain, despite the statistically significant reduction in its motility (Fig. 2a, b). Furthermore, we observed no differences in other aspects of rod-cell morphology, including length, width, aspect ratio, cell area, curvature of the longitudinal axis of cells, overall cell outline curvature, and maximum curvature of cell poles (Fig. 2c-i, respectively). We conclude that altered overall swimming speed through *cetZ* modification does not always correlate with measurable changes in cell morphology.

### Interdependency and GTPase requirements of *cetZ1* and *cetZ2* phenotypes

To identify any potential coordination of CetZ1 and CetZ2 functions leading to the above phenotypes, we also investigated the dependency of the *cetZ1/2* overexpression phenotypes on the presence of *cetZ1* or *cetZ2* (Fig. S1, S2). Overexpression of *cetZ1* in the Δ*cetZ2* background reduced motility compared to the control, like it did in the wildtype background. On the other hand, overexpression of *cetZ2* in the Δ*cetZ1* background failed to rescue the Δ*cetZ1* motility defect and did not clearly stimulate motility. Thus, the mild stimulation of motility by *cetZ2* depends on the presence of *cetZ1*, whereas the inhibition of motility by *cetZ1* overexpression does not depend on *cetZ2*.

To test the importance of the expected GTPase activity of CetZs, we overexpressed GTPase active-site mutants (*cetZ1*.E218A or *cetZ2*.E212A) in the wildtype (H26) background. Previous work showed that expression of these mutants hyper-stabilizes the protein assemblies *in vivo* and results in a heterogeneous population of irregularly shaped cells in the *H. volcanii* H98 background (Duggin et al., 2015). Consistent with these, we found that *cetZ1*.E218A or *cetZ2*.E212A strongly inhibited motility and rod shape in both the H26 wildtype and respective Δ*cetZ1* or Δ*cetZ2* backgrounds (Fig. S1, 2, 3). These results confirm the dominant-inhibitory behaviour of these mutants and suggest that GTPase activity is necessary for the overexpression phenotypes of *cetZ1* and *cetZ2*.

### CetZ1 and CetZ2 influence the number of cell surface filaments

Given that the *cetZ1/2* overexpression results above suggested that CetZs might affect motility in both shape-dependent and shape-independent ways, we began investigating the hypothesis that CetZs influence the assembly or function of other cell envelope or motility structures such as the archaellum or chemosensory arrays. To determine whether CetZs influence the overall appearance of surface filaments, such as pili and archaella, we initially used cryo-EM of live cells (e.g., Fig. 3a) to compare the wildtype, *cetZ1* deletion, and *cetZ2* overexpression strains, which showed the most substantial and opposing effects on motility (Fig. 1). Deletion of *cetZ1* resulted in a significant reduction in the number of surface filaments per cell, whereas overexpression of *cetZ2* resulted in slightly more (Fig. 3b). As deletion of *cetZ1* or overexpression of *cetZ2* did not affect cell adhesion (Fig. S4), the differences in the number of surface filaments probably reflects the number of archaella rather than adhesive pili, consistent with the observed motility phenotypes of these strains.

**Figure 3.**
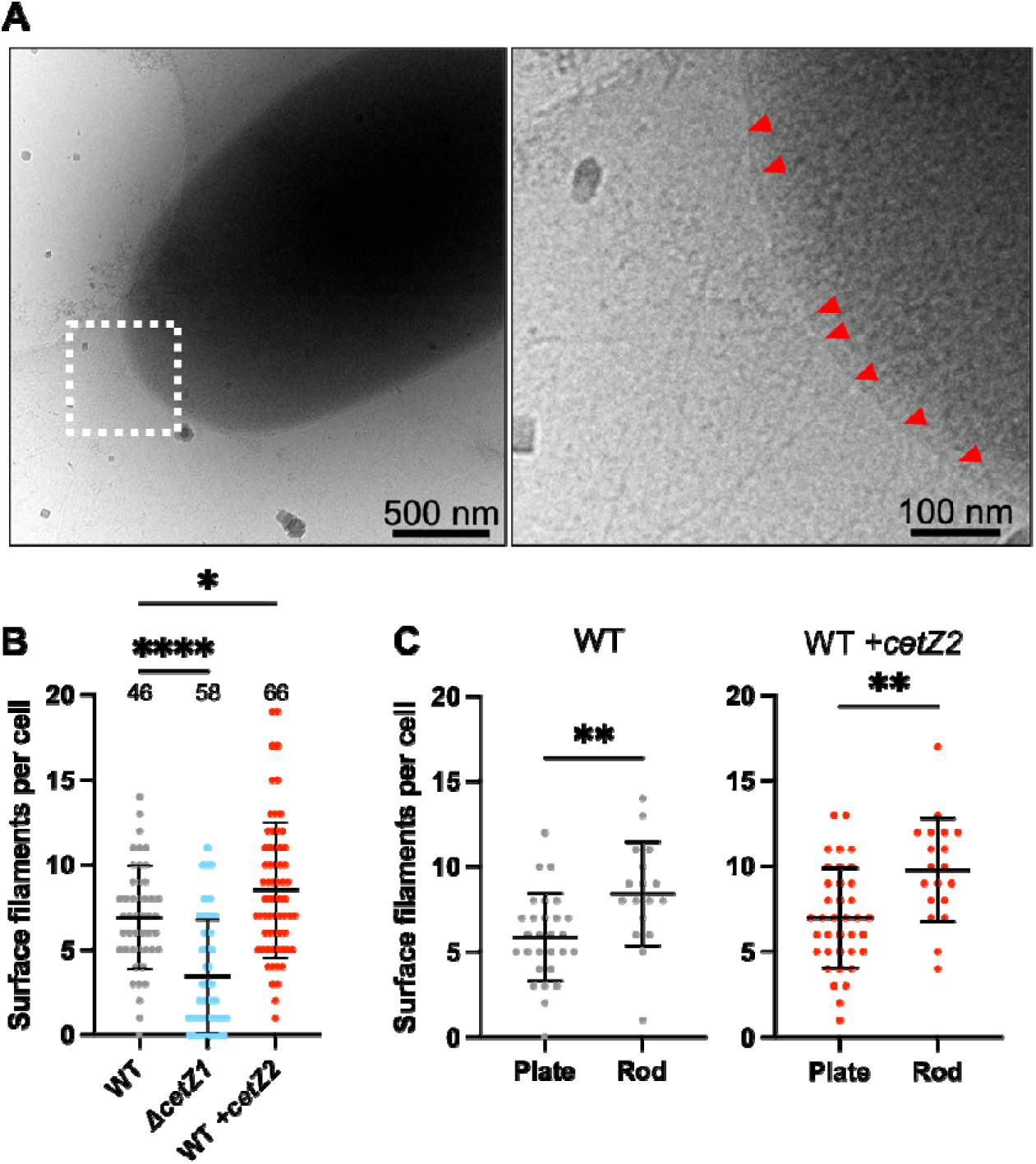
Surface filament assembly in motile cells and *cetZ* mutants. A) Representative CryoEM image of H26 wildtype cells (left) and zoom (right) showing surface filaments (red tick marks). **B)** The number of surface filaments per cell were manually counted using images obtained in **A** for H26 wildtype, Δ*cetZ1*, and WT +*cetZ2* strains. The number of cells analysed per strain is indicated above the data for each strain. **C)** For rod forming strains, WT and +*cetZ2*, the number of surface filaments per cell was determined for rod and plate-shaped cells. Data in **B** and **C** was obtained from the same sample and show mean and standard deviation. **B** uses one-way ANOVA as a statistical test, and **C** uses an un-paired T-test. ****p<0.0001, **p<0.002, *p<0.02.

We also compared the number of surface filaments in rod- and plate-shaped cells in each sample. This showed that rod-shaped cells had significantly more surface filaments compared to plate-shaped cells in both the wildtype and *cetZ2* overexpression strains (Fig. 3c). Interestingly, the number of surface filaments in the Δ*cetZ1* strain was still lower than the number of surface filaments in plate-shaped wildtype cells (Fig. 3b, c). Plate cells can be motile (including Δ*cetZ1* cells), albeit with reduced speeds (Fig. 1d) (Duggin et al., 2015). These observations suggest that the motility defect of Δ*cetZ1* is not only associated with its rod shape defect, but also a reduction in the appearance of cell surface filaments.

### Functional CetZ-mTq2 fusions localise to cell poles in motile cells

CetZ1 was reported to localise strongly to cell poles in motile rod-shaped cells (Duggin et al., 2015). However, since the CetZ1-GFP fusion was not fully functional as a sole copy in rod-development, it remained unclear whether the polar localization reflected a native function of CetZ1. Subsequent studies of CetZ1 localization using an almost fully functional CetZ1-mTurquoise2 (mTq2) fusion have not investigated CetZ1 localization in motile cells (de Silva et al., 2024; Ithurbide et al., 2024). Here, we used the existing CetZ1-mTq2 fusion and generated a CetZ2-mTq2 fusion to observe their localization in motile cells.

CetZ1-mTq2 and CetZ2-mTq2 were first tested for their capacity to support motility in the wildtype background and their respective *cetZ* knockout backgrounds (Fig. 4a), which resulted in motility halos that were on average 71% and 107% of the size of those produced by expression of untagged *cetZ1* and *cetZ2*, respectively; these differences in motility were statistically significant between Δ*cetZ1* +*cetZ1* and Δ*cetZ1* +CetZ1-mTq2, but not between Δ*cetZ2* +*cetZ2* and Δ*cetZ2* +CetZ2-mTq2. CetZ1-mTq2 and CetZ2-mTq2 both slightly reduced the circularity of motile rods when expressed in either the wildtype or knockout backgrounds (Fig. 4b), but both fusion proteins were produced at similar levels to their untagged counterparts (Fig. S5).

**Figure 4.**
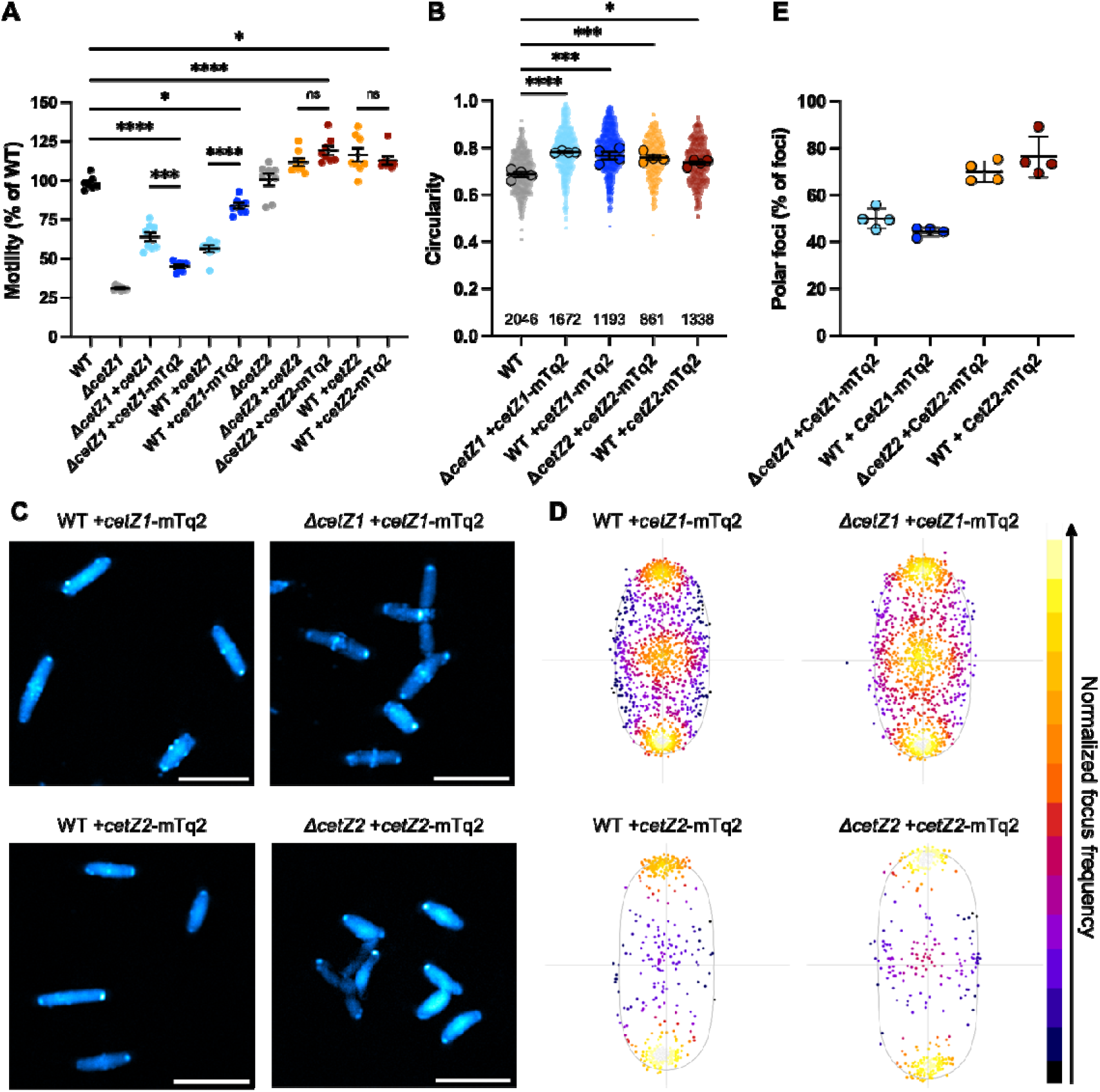
Localization of CetZ1 and CetZ2 in motile cells. CetZ1-mTq2 and CetZ2-mTq2 fusion proteins were produced in H26 wildtype cells and their respective knockout backgrounds. H26 wildtype, Δ*cetZ1,* and Δ*cetZ2* with pTA962, H26 *+cetZ1,* H26 *+cetZ2*, Δ*cetZ1* +*cetZ1,* and Δ*cetZ2* + *cetZ2* were used as controls. **A)** Strains were inoculated onto Hv-Cab soft-agar with 1 mM L-Tryptophan and their motility halo diameters were quantified. Individual points represent independent replicates. **(B)** Cells expressing *cetZ1*-mTq2 or *cetZ2*-mTq2 were extracted from the edge of the motility halo and their circularity was measured from phase-contrast microscopy images (not shown). Small data points represent individual cells, while large data points represent the mean of all cells measured from one replicate culture. Error bars indicate mean and standard error. The number of cells (n_c_) pooled from all independent replicates (n_r_) for each strain are indicated below the data for each strain and apply to all panels. **C)** Fluorescence microscopy was also conducted to observe CetZ1 and CetZ2 localization in these cells. **D)** Heatmap representations of the position of CetZ1-mTq2 and CetZ2-mTq2 foci in motile cells. Normalized cell aspect ratio was used to generate a typical cell shape of the population for mapping foci. **E)** Proportion of detected CetZ foci located at cell poles. In **A** and **E**, mean and standard deviation are shown. One-way ANOVA was used as a statistical test, and only significant and/or relevant comparisons are shown on graphs. ****p<0.0001, ***p<0.0002, ns=not significant.

In motile cells extracted from the halo edge, CetZ1-mTq2 and CetZ2-mTq2 displayed bright foci or patches at the cell poles (Fig. 4c-e). CetZ1-mTq2 showed additional patchy localization around the envelope and at mid-cell (Fig. 4c), like previously described (de Silva et al., 2024; Duggin et al., 2015; Ithurbide et al., 2024). These similarities and differences in CetZ1-mTq2 and CetZ2-mTq2 foci were clearly reflected in heatmap representations of foci position within a schematic representation of cell shape of the population, based on normalized longitudinal and lateral cell axes (Fig. 4d). No differences in localization were observed when CetZ1-mTq2 and CetZ2-mTq2 were produced in the either the wildtype or their respective knockout background. These results confirm that the CetZs strongly localize to the cell poles in mature motile cells and might carry out some of their specific functions there.

### CetZ1 strongly promotes the assembly of the archaellum base protein ArlD1

Given the above results, we sought to determine whether deletion of *cetZ1* or *cetZ2* or overexpression of the wildtype or GTPase mutants affect the assembly or positioning of key motility marker proteins at the subcellular level. These included ArlD1 as a marker for the archaellum base (Fig. 5), CheW1 as a marker for chemosensory arrays (Fig. 6), and CheY (Fig. 7), which is the effector protein that shuttles between chemosensory arrays and the archaellum base. Their localization in motile cells have been previously well characterised (Li et al., 2019; Li et al., 2020) and expression of ArlD1-GFP, GFP-CheW1, and CheY-GFP did not cause motility or cell shape changes (Fig. S6a-f).

**Figure 5.**
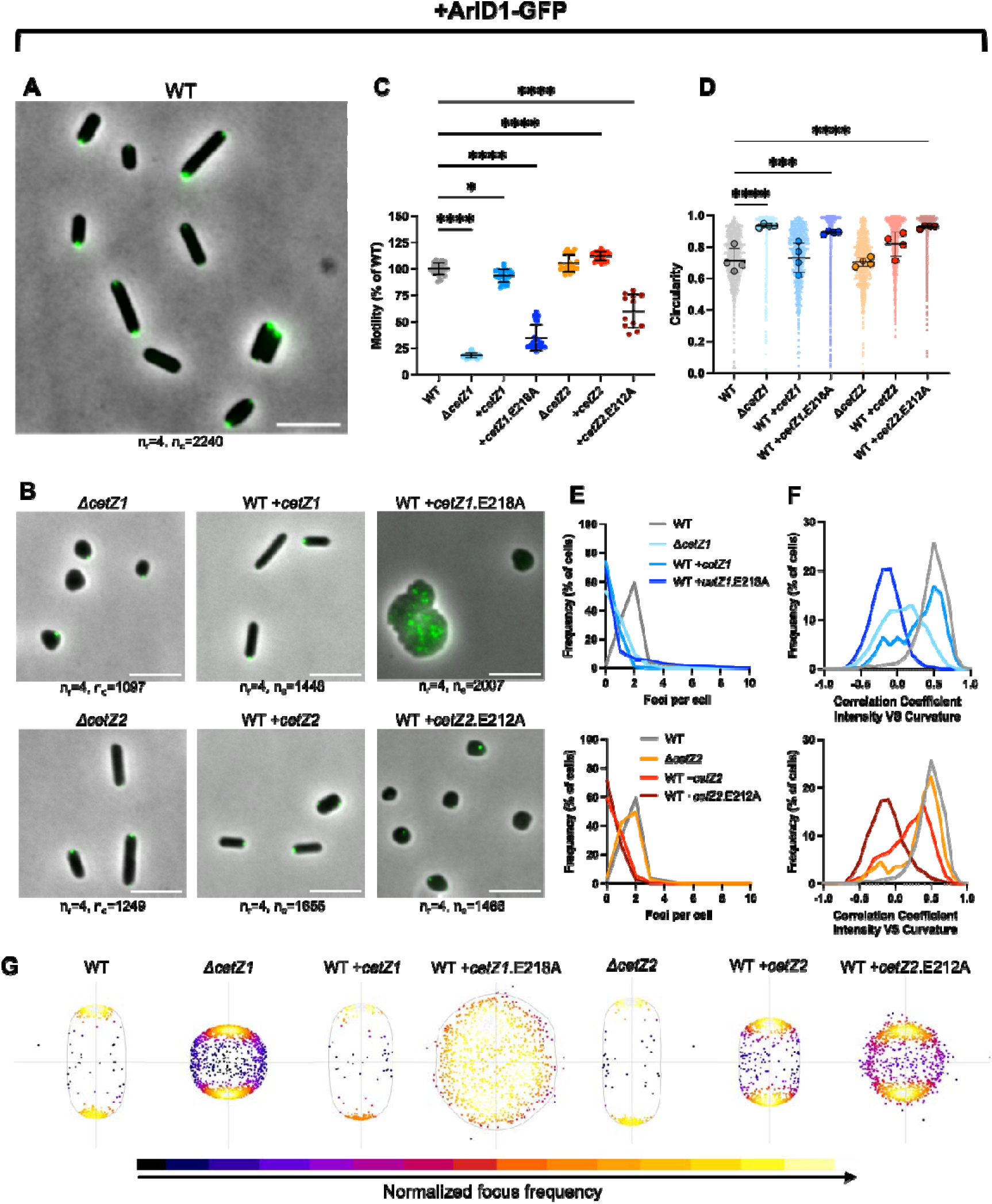
ArlD1-GFP localization in *cetZ1/2* mutant strains. ArlD1-GFP was produced from pSVA3919 in H26 wildtype, Δ*cetZ1*, and Δ*cetZ2*, or was co-expressed with *cetZ1*, *cetZ1*.E218A, *cetZ2*, or *cetZ2*.E212A from pHJB31, pHJB34, pHJB37, or pHJB40, respectively, in H26 wildtype. These strains were inoculated onto Hv-Cab soft-agar with 1 mM L-Tryptophan and extracted from the halo-edge for phase-contrast and fluorescence microscopy **(A, B)**. Scale bar: 5 μm. the number of cells (n_c_) pooled from all independent replicates (n_r_) for each strain are indicated below images and apply to all panels. **C)** Motility halo diameters for each strain were quantified. Individual points represent independent replicates. Mean and standard deviation is shown. **D)** Phase-contrast images were used to measure circularity. Large data points represent the mean of all cell circularities in an independent replicate, and small data points represent individual cells. Mean and standard deviation of replicate means are shown. One-way ANOVA was used as a statistical test (comparing replicate means in **D**). ****p<0.0001, ***p<0.0002, **p<0.002, *p<0.02. Only significant comparisons are shown. Circularity varied somewhat between replicate halo samples of the same strain (Fig. S8d), which is likely due to differences in the sampling of some plate shaped cells that are present just inside the outer edge of the halo in soft agar (Duggin et al., 2015). **E)** Frequency distributions of the number of foci per cell in each strain, and **F)** the Pearson Correlation Coefficient between local membrane curvature and fluorescence intensity. In **E** and **F**, data for WT +ArlD1-GFP is displayed on all graphs for reference, and the key refers to both panels. **G)** Heatmap representations of ArlD1-GFP foci frequency with respect to the average cell shape normalized by the two major cell axes, in the indicated strain backgrounds.

**Figure 6.**
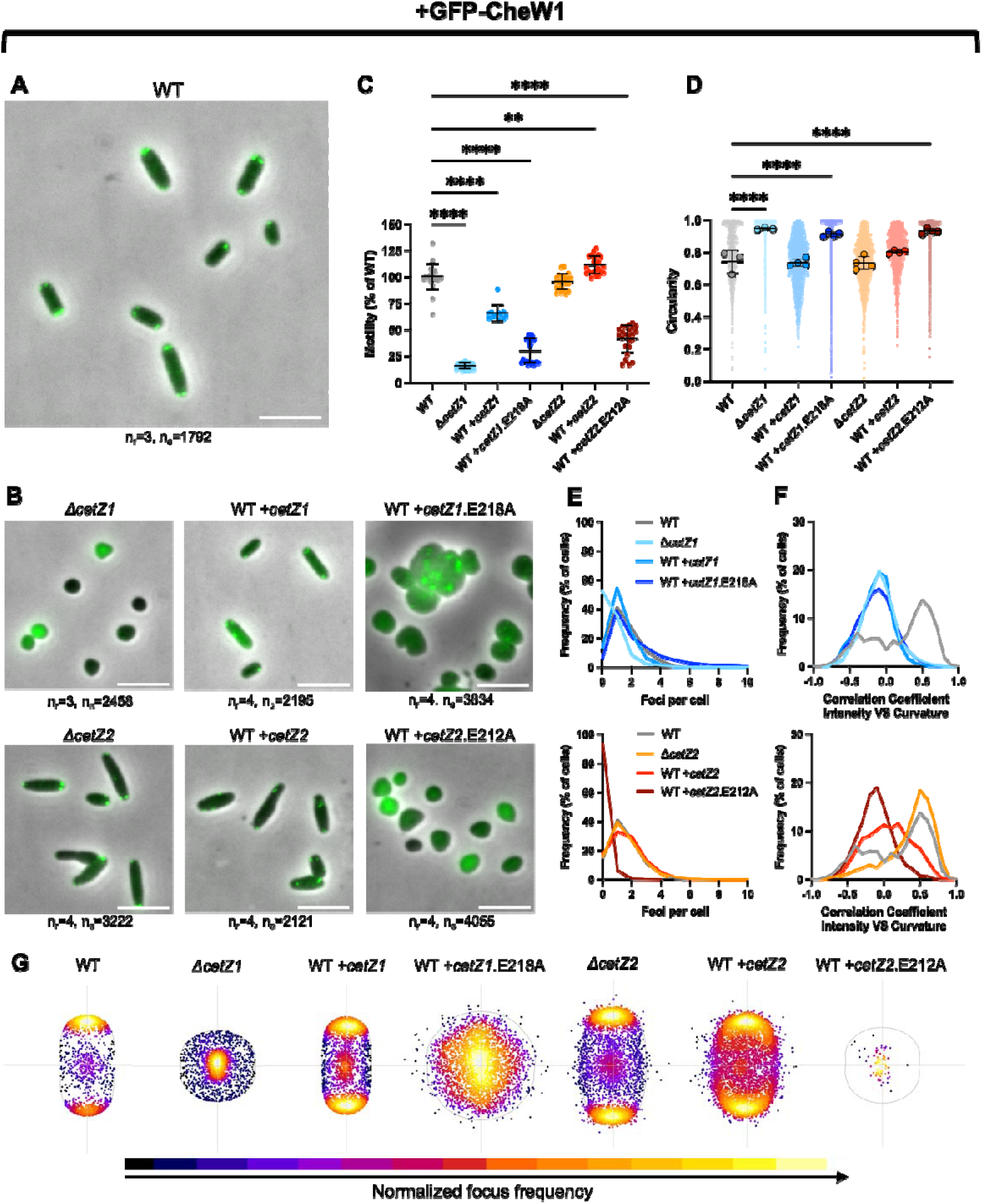
GFP-CheW1 localization in *cetZ1/2* mutant strains. GFP-CheW1 was produced from pSVA5031 in H26 wildtype, Δ*cetZ1*, and Δ*cetZ2*, or was co-expressed with *cetZ1*, *cetZ1*.E218A, *cetZ2*, or *cetZ2*.E212A from pHJB32, pHJB35, pHJB38, or pHJB41, respectively, in H26 wildtype. These strains were inoculated onto Hv-Cab soft-agar with 1 mM L-Tryptophan and extracted from the halo-edge. **A-G** is as described in Figure 5.

**Figure 7.**
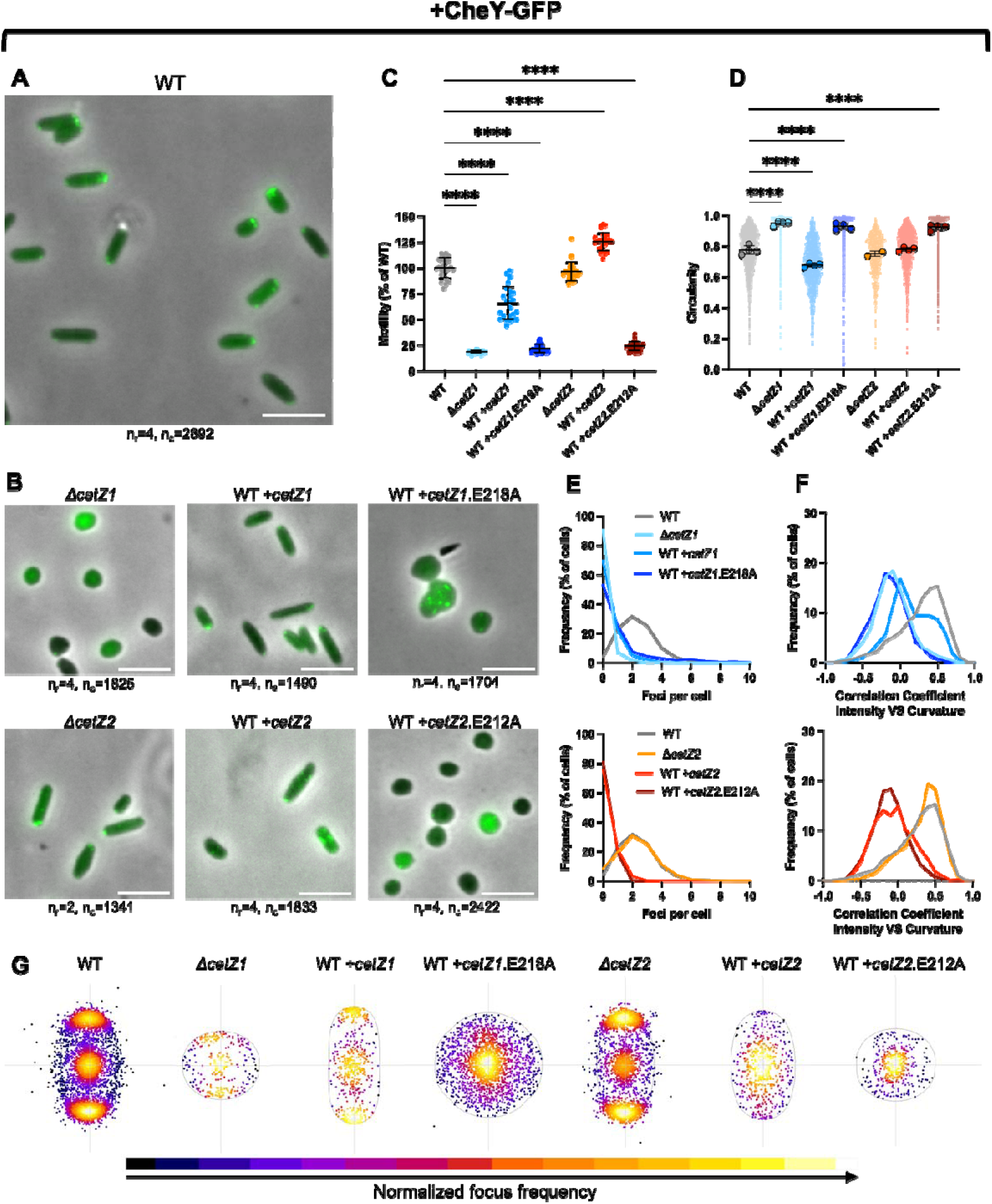
CheY-GFP localization in *cetZ1/2* mutant strains. CheY-GFP was produced from pSVA5611 in H26 wildtype, Δ*cetZ1*, and Δ*cetZ2*, or was co-expressed with *cetZ1*, *cetZ1*.E218A, *cetZ2*, or *cetZ2*.E212A from pHJB33, pHJB36, pHJB39, or pHJB42, respectively, in H26 wildtype. These strains were inoculated onto Hv-Cab soft-agar with 1 mM L-Tryptophan and extracted from the halo-edge. **A-G** is as described in Figure 5.

In wildtype motile cells, ArlD1-GFP localised as one or most commonly two foci of variable intensities at the cell poles (Fig. 5a, 5e, 5g), like previously observed (Li et al., 2019). Cells with two or more foci tended to be more elongated than the single- or zero-focus cells (Fig. S6g). The great majority of detected ArlD1-GFP foci were located at the cell poles (>97%, Fig. 5a, 5g, and S7a). Consistent with this, the local curvature of the cell outline strongly correlated with fluorescence (Fig. 5f, upper graph; Fig. S8 illustrates the analysis approach).

In the Δ*cetZ1* background, most cells showed no ArlD1-GFP localization or only one focus adjacent to the edge of the cell (Fig.5b, 5e, 5g). These cells showed the expected loss of rod shape (Fig. 5b, 5d, S9a), and there was no clear correlation on average between local cell curvature and ArlD1-GFP localization around the cell outline (Fig. 5f). However, the foci position heatmap indicated a moderate tendency of the detected foci to be located at the ends of the slightly longer of the two cell axes (Fig. 5g).

When *cetZ1* was overexpressed, motile cells showed the normal rod morphology (Fig. 5b, 5d), including normal cell width and curvature of the cell outline and the poles (Fig. S9a-c). Importantly, the number of ArlD1-GFP foci (Fig. 5e) and the average correlation of fluorescence with cell curvature (Fig. 5f) were both significantly reduced compared to the wildtype background. These defects in assembly and positioning of ArlD, despite the normal rod shape, are thus consistent with the reduced motility caused by *cetZ1* overexpression (Fig. 5c).

Given that the *cetZ* GTPase active-site mutants localize aberrantly and deform the cells (Duggin et al., 2015), we also investigated their effects on ArlD localization. At relatively high overexpression, *cetZ1*.E218A can cause moderate cell enlargement, signifying an interference with cell division (e.g., Fig. 5b, S2a); we noticed that some cells were substantially enlarged, in which ArlD1-GFP appeared as many foci distributed throughout the cell and not at the cell edges (Fig. 5b). However, most cells were smaller and showed low or no localized fluorescence (Fig. 5b, 5e, 5g). Consistent with these observations, there was a weak negative correlation on average between ArlD1-GFP fluorescence and curvature of the cell outline (Fig. 5f). Overexpression of *cetZ1*.E218A therefore strongly interferes with both the assembly and positioning of ArlD1-GFP foci in *H. volcanii*.

### cetZ2 overexpression influences the assembly of ArlD1

Deletion of *cetZ2* (Fig. 5b) had no detected effect on the assembly (Fig. 5e) or positioning of ArlD1-GFP foci at the cell poles (Fig. 5f, 5g, S7a), consistent with the normal motility and cell shape phenotypes exhibited by Δ*cetZ2* (e.g., Fig. 1, Fig. 2) (Duggin et al., 2015). When *cetZ2* was overexpressed, ArlD1-GFP assembly at cell poles appeared moderately reduced (Fig. 5b, 5f, 5g, S7a) and showed fewer foci per cell (Fig. 5e), despite its slightly elevated motility and number of surface filaments (Fig. 1a, 1b, 3b). A possible explanation is that the increased number of surface filaments (presumably archaella) were not reflected in the ArlD1-GFP imaging due to close clustering of archaella at the poles.

Expression of the *cetZ2*.E212A GTPase mutant resulted in a milder cell shape effect than *cetZ1*.E218A (Duggin et al., 2015), where rod shape was inhibited but the plate-like cells were not highly irregular or enlarged (e.g. Fig. 5b, S2). Under these conditions, ArlD1-GFP appeared as infrequent foci that were often not associated with the cell edge (Fig. 5b, 5e) and showed a slight negative correlation with cell curvature on average (Fig. 5f). Thus, like *cetZ1*.E218A, overexpression of *cetZ2*.E212A strongly interferes with the positioning of ArlD1-GFP.

Since *cetZ2* deletion showed no impact on ArlD1-GFP positioning or assembly, these results suggest that *cetZ2* is not substantially involved in archaellum assembly or motile rod development under the standard conditions investigated. However, overexpression of *cetZ2* appears to mildly stimulate aspects of motility via an unknown mechanism requiring *cetZ1*, which appears to be accompanied by changes in archaellum positioning and number.

### Assembly of the chemotaxis array protein CheW1 is highly dependent on CetZ1

Another major component of motility is the chemotaxis cell surface array that senses and conveys extracellular signals to the archaellum to control the overall direction of swimming. In the wildtype background, we confirmed that the array marker protein GFP-CheW1 localized to polar and subpolar regions as multiple foci or patches/filaments (Fig. 6a), as expected (Li et al., 2019). Most of the cells were rods, with a small fraction of plate-shaped cells in these samples (Fig. 6a, 6d), and we observed that cells with more foci tended to show greater elongation (Fig. S6h).

Strikingly, deletion of *cetZ1* resulted in a diffuse signal for GFP-CheW1, with only a very small proportion of cells displaying distinct faint foci (Fig. 6b,6e) compared to the typically 1-3 or more bright polar/subpolar foci in the wildtype background (Fig. 6a, 6e, 6g). In some cells, a broad GFP-CheW1 zone appeared around the middle of the cell (Fig. 6b), which was often detected as a focus (Fig. 6e, 6g), however it clearly does not resemble the sharply resolved foci of normal chemotaxis arrays. Consistent with this, the fluorescence intensity of GFP-CheW1 around the cell outline in Δ*cetZ1* also was not correlated with cell curvature (Fig. 6f).

Overexpression of *cetZ1* did not have a substantial impact on the number of GFP-CheW1 foci per cell (Fig. 6e), but it did have a moderate influence on the position of foci in the cell (Fig. 6g) and their localization at areas of cell curvature (Fig. 6f, S7b), which is consistent with the moderate reduction in motility (Fig. 6c). The *cetZ1*.E218A GTPase mutant had a similar inhibitory effect on GFP-CheW1 localization as it did on ArlD1-GFP, substantially disrupting cell shape, GFP-CheW1 localization (Fig. 6b, 6g), and its association with cell curvature (Fig. 6f). Together these observations suggest CetZ1 is a key driver of CheW1 assembly.

Deletion of *cetZ2* had no detected effect on foci number (Fig. 6e) or positioning of GFP-CheW1 (Fig. 6f, 6g). Although overexpression of *cetZ2* caused no clear difference in GFP-CheW1 foci number (Fig. 6e) it resulted in a reduced average correlation of GFP-CheW1 fluorescence with curvature (Fig. 6f) and, consistent with this, a moderate redistribution of some of the foci throughout the cells (Fig. 6g). In contrast, expression of *cetZ2*.E212A resulted in a diffuse signal for GFP-CheW1 (Fig. 6b), with the great majority of cells having no detectable GFP-CheW1 foci (Fig. 6e) and there was essentially no clear correlation between fluorescence and cell curvature (Fig. 6f, 6g), similar to the effects of the *cetZ1*.E218A mutant. These results suggest that *cetZ2* is not directly required for the normal assembly of chemotaxis arrays but has the capacity to disrupt their formation when overexpressed, particularly in the GTPase-inactive form that also has dominant effects on cell shape (Fig. 6b).

### CheY assembly is highly dependent on CetZ1

We next sought to determine whether CetZs also influence the effector protein, CheY (Fig. 7a), which shuttles between chemosensory arrays and the archaellum base as part of the signal transduction pathway which results in switching of rotation direction of the archaellum. Deletion of *cetZ1* resulted in mainly diffuse localization of CheY-GFP, with some distinct foci at the cell edge remaining (Fig. 7b, 7g). Overexpression of *cetZ1* did not change the localization pattern of CheY-GFP, but it did reduce the frequency of foci (Fig. 7b, 7e-g). As with ArlD1-GFP and GFP-CheW1, expression of *cetZ1*.E218A also produced a sub-population of large and highly irregular cell shapes with multiple CheY-GFP foci (Fig. 7b), but in most cells, CheY-GFP failed to form foci (Fig. 7e).

Deletion of *cetZ2* had no detected effect on assembly or positioning of CheY-GFP (Fig. 7b, S7c). Overexpression of *cetZ2* resulted in fewer CheY-GFP foci per cell, and like GFP-CheW1, the correlation between CheY-GFP localization and cell curvature was reduced. Expression of *cetZ2*.E212A had similar effects as it did on CheW1 assembly (Fig. 7b, 7e-g). The CheY-GFP signal was diffuse or weakly centrally located in most of these cells (Fig. 7b, 7g).

Overall, the results show a similar pattern of behaviour amongst the archaellum and chemotaxis marker proteins in response to the various *cetZ* mutant backgrounds, strongly implicating cetZ1 as a key driver of both the assembly and correct polar positioning of the motility machinery in *H. volcanii*, most likely via a mechanism that is independent of the rod-shape of the motile cells.

### Dual localization studies of CetZ1 and motility marker proteins

Given the above findings, we sought to further investigate the relationships between ArlD1, CheW1, and CheY assembly and CetZ1 by assessing the extent of polar co-localization between the motility markers and CetZ1 (Fig. 8a-c). When a functional CetZ1-mCh fusion (Ithurbide et al., 2024) was produced in tandem with either ArlD1-GFP, GFP-CheW1, or CheY-GFP, all four fusions displayed localization patterns in swimming cells equivalent to those observed when they were expressed individually (Fig. 4c, 5a, 6a, 7a, respectively). To determine the degree to which ArlD1-GFP foci co-occur with CetZ1-mCh foci, each ArlD1-GFP focus was identified and then categorized into one of three patterns with respect to CetZ1-mCh foci (Fig. 8d). This revealed that the majority (73%) of ArlD1-GFP localizations overlapped with a polar focus of CetZ1-mCh (Fig. 8d(ii)), 26% of ArlD1-GFP foci had no detected CetZ1-mCh focus at the same pole (Fig. 8d(i)), and less than 1% were separated from a CetZ1-mCh focus at the same pole (Fig. 8d(iii)). We also counted cell poles with either one, or both focus types present (irrespective of their degree of overlap) and found that 13.8% of poles had both ArlD1-GFP and CetZ1-mCh, which was significantly higher than the 8.1% that would be expected based on a random allocation of the individual foci to each pole (Fig. 8g).

**Figure 8.**
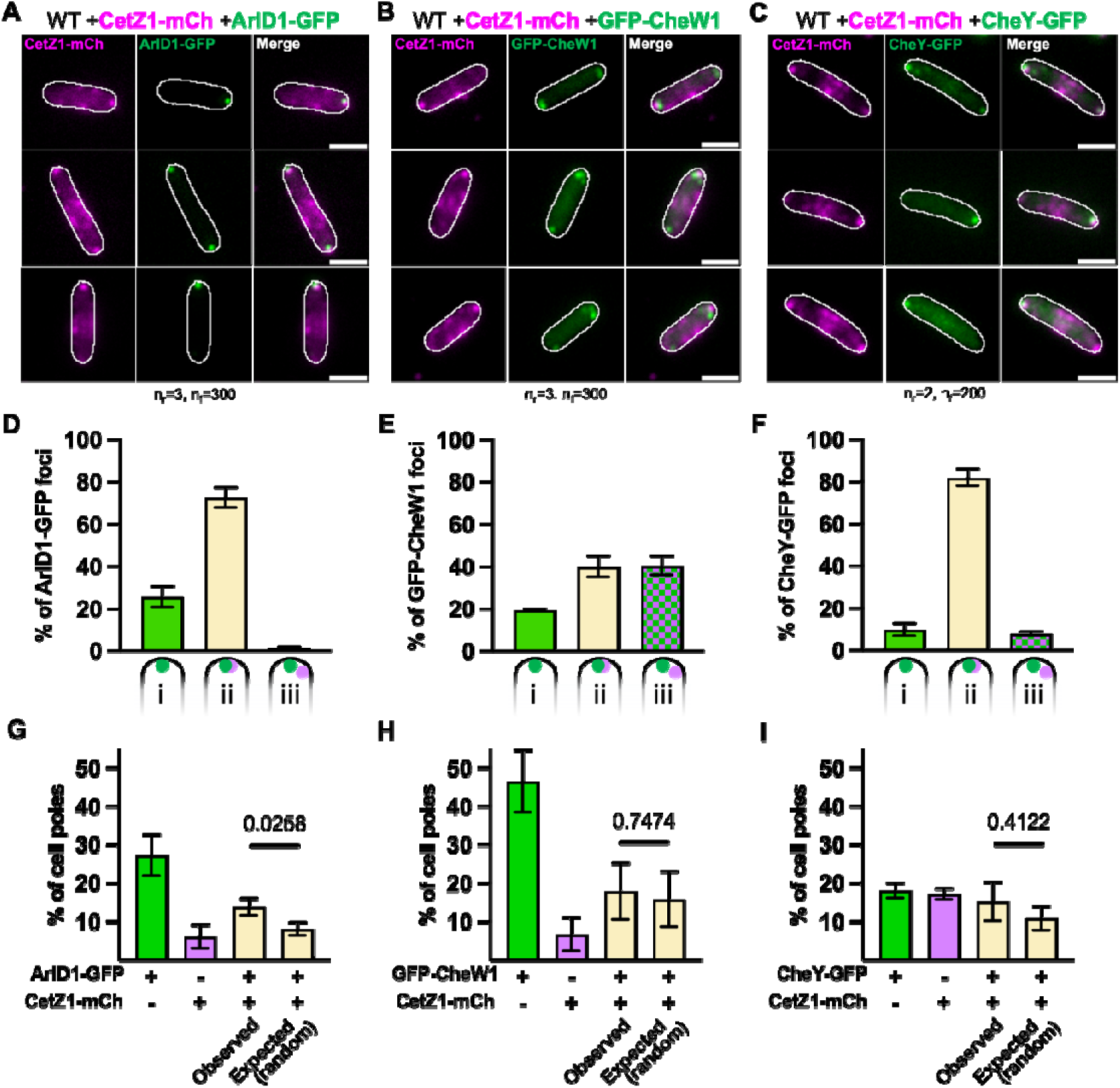
Dual localization of CetZ1-mCherry with ArlD1, CheW1 and CheY GFP fusions. **A)** ArlD1-GFP, **B)** GFP-CheW1, or **C)** CheY-GFP were produced in tandem with CetZ1-mCh in H26 wildtype cells from plasmids pHJB69, pHJB70, and pHJB71, respectively. Cells were inoculated onto Hv-Cab soft-agar with 1 mM L-Tryptophan and extracted from the halo edge for fluorescence microscopy. At least two culture replicates were carried out for each strain. To assess the degree of CetZ1-mCh localisation at cell poles with a localised motility marker, 100 foci of motility marker proteins were first identified without viewing CetZ1-mCh localization for each culture replicate. The total number of foci (n_f_) from independent replicates (n_r_) are indicated below panels **A**, **B**, and **C**, and apply to **D** and **G**, **E** and **H**, and **F** and **I** respectively. The selected foci of **D)** ArlD1-GFP, **E)** GFP-CheW1, and **F)** CheY-GFP were classed in one of three categories: (i) no focus of CetZ1-mCh detected at the same pole, (ii) overlapping (fully or partially) with CetZ1-mCh focus; or (iii) not overlapping with a CetZ1-mCh focus detected at the same cell pole. The presence or absence of CetZ1-mCh, **G)** ArlD1-GFP, **H)** GFP-CheW1, and **I)** CheY-GFP was scored at each cell pole across the entire population. This was conducted separately for each independent (culture) replicate, and presence/absence data is represented as a percentage of cell poles. ‘+’ indicates present, ‘-’ indicates absent. **D-I)** show mean and standard deviation. A two-tailed T-test was conducted to statistically compare the observed and expected proportions of cell poles with both the motility marker and CetZ1-mCh. The resulting p-values are shown.

Foci co-localization analysis of CetZ1-mCh with GFP-CheW1 (Fig. 8b, e, h) demonstrated that polar CetZ1-mCh foci were less closely associated with GFP-CheW1 foci positioning compared to ArlD-GFP; 40.5% of GFP-CheW1 foci did not overlap with CetZ1-mCh foci, even when located at the same cell pole (Fig. 8e(iii)), and only 40% of GFP-CheW1 foci overlapped with CetZ1-mCh foci (Fig 8e(ii)), and the remaining ∼20% of GFP-CheW1 foci showed no CetZ1-mCh focus at the same pole (Fig 8e(i)). This poorer co-localization than with ArlD1-GFP (Fig. 8d, 8e) is consistent with the differing localization patterns of the chemotaxis arrays (often subpolar) and archaella (polar) (Fig. 6, 7). Finally, analysis of CheY-GFP foci showed that 82% had an overlapping CetZ1-mCh focus and only 10% had no CetZ1-mCh focus at the same pole (Fig. 8c, 8). Both CheW-GFP and CheY-GFP did not show a significant propensity to co-occur with CetZ1-mCh at poles compared to a random allocation (Fig. 8h, i).

Overall, these co-localization patterns between CetZ1 and motility proteins appeared less intimate than what would be expected for an obligate or stoichiometric co-assembly of CetZ1 with the motility markers. Rather, their imperfect overlap and co-incidence in polar regions suggested that CetZ1 may have a structural or organisational role at the cell pole that indirectly influences the correct assembly and positioning of the motility machinery.

## Discussion

Archaeal CetZ tubulin-like proteins have previously been implicated in the control of cell shape and swimming motility (Duggin et al., 2015). In *H. volcanii*, development of rod-shaped cell types requires CetZ1 (de Silva et al., 2024; Duggin et al., 2015) and this morphotype is highly correlated with swimming motility (Brown and Duggin, 2023; Duggin et al., 2015; Schwarzer et al., 2021).

Here, we found that although some manipulations of CetZs (i.e., Δ*cetZ1*, overexpression of *cetZ2*, and expression of the dominant GTPase-defective mutants *cetZ1*.E218A or *cetZ2*.E212A) exhibited defects in both rod-shape and overall swimming speed, overexpression of wildtype *cetZ1* resulted in reduced overall swimming speed without affecting the consistent rod cell shape of the swimming cells. Previous work showed that *cetZ1* overexpression does not always cause phenotypic defects, because it stimulated the generation of rod morphologies in liquid cultures that otherwise show mainly plate cell morphologies (Duggin et al., 2015). These observations suggest that, in addition to their role in promoting rod cell shape to optimize motility, CetZs might contribute to motility in other ways too.

Three additional observations are consistent with the existence of CetZ functions that improve swimming beyond the basic hydrodynamic streamlining afforded by the rod cell morphology. Firstly, we found that a functional CetZ1 fluorescent protein fusion strongly localises to the poles of actively motile rods, which have already fully completed the rod morphogenesis process. This polar localization differs substantially compared to the longitudinal filaments and dynamic CetZ1-FP foci seen during rod formation and in rods from liquid culture (de Silva et al., 2024; Duggin et al., 2015). It also appears to be regulated by the MinD2/4 proteins (Brown and Duggin, 2024). These observations are consistent with an additional role for CetZ1 specifically at the poles of motile rods. Secondly, the Δ*cetZ1* strain showed fewer cell surface filaments (likely archaella) than wildtype plate-shaped cells, which are the same shape as Δ*cetZ1* cells. Thirdly, overexpression of *cetZ2* resulted in slightly greater overall swimming speed and cell surface filament numbers compared to the wildtype, yet also showed a minor reduction in the degree of rod cell morphology. These findings suggest that the cell shape is not the only factor controlled by *cetZ* that influences the assembly of the motility machinery. Based on these initial observations, we sought to address the hypothesis that CetZs contribute to motility by promoting the polar assembly of archaella and chemosensory array protein complexes.

Deletion of *cetZ1* or overexpression of *cetZ1* or *cetZ1*.E218A all substantially reduced the assembly and polar positioning of the archaellum and chemotaxis marker proteins, ArlD1, CheW1, and CheY. The effects were of varying degrees that were approximately consistent with the magnitude of their motility defects. Importantly, deletion of *cetZ1* caused major defects in the frequency of assembly of the motility marker proteins, and their remaining localization patterns differed substantially, suggesting that the effects of CetZ1 on ArlD1, CheW1, and CheY localization are not only related in a simple way to cell shape. Furthermore, *cetZ1* overexpression interfered with the proper assembly of the motility proteins without changing the rod shape of motile cells. It also consistently reduced motility in all of the wildtype and motility marker backgrounds. Such a negative response of a phenotype to overexpression may seem counterintuitive at first but is not uncommon and may result from steric interference or concentration-dependent effects on protein function (Prelich, 2012). Excess CetZ1 might thus cause alterations in molecular assemblies at the cell pole, without altering morphology at a cellular level, and thereby lead to misplacement or failure in the proper assembly of the motility machineries.

While our results do not completely rule out undetected subtle effects of some of the *cetZ1* perturbations on rod cell shape development that substantially influence motility machinery localization, we believe the simplest explanation for our combined results is that the polar-localized CetZ1 locally promotes the assembly of the motility machinery. This would represent an additional role to CetZ1’s function in rod cell morphogenesis, which is also likely to contribute to motility through streamlined hydrodynamics and establishment of cell polarity (Duggin et al., 2015). A role for CetZ1 in motility that is not directly related to rod shape may be further supported by the observation that loss of polar-localised CetZ1 in the absence of *minD2* is correlated with decreased motility but no observed impact on rod shape (Brown and Duggin, 2024).

The potential involvement of CetZ2 in motility is less clear than for CetZ1. Deletion of *cetZ2* caused no detected defects in motility or assembly of the motility machinery. Although overexpression of *cetZ2* resulted in a mild increase in motility and the number of cell surface filaments, the increase in motility was subtle and required the presence of c*etZ1*. Despite this, *cetZ2* overexpression also resulted in somewhat fewer polar foci of the motility marker proteins. One potential explanation is that *cetZ2* overexpression synthetically promotes clustering of the motility proteins at the cell pole, which are not resolved by fluorescence microscopy. Thus, whether CetZ2 has a natural role in *H. volcanii* motility in standard or other conditions is currently unclear.

How might CetZ1 contribute to the structure of the cell poles and assembly of the motility machinery there? Our dual-localization studies indicated that polar foci of CetZ1 and the motility marker proteins often overlapped but not always. In particular, the archaellum marker, ArlD1-GFP, and CetZ1-mCh appeared to have a specific propensity to co-localize at cell poles. However, the imperfect co-localization and partial independence of localization suggests that none of the motility machinery markers form an obligate or stoichiometric co-assembly with CetZ1. Since CetZ1 has a strong impact on assembly of all the motility markers—particularly CheW1, despite showing the lowest polar co-localization with CetZ1—our results suggest that CetZ1 probably has a distinct function in establishing the structure of the cell pole, which in turn plays a significant role in the proper assembly of the motility machineries.

Polar cap-like structures have been identified in several archaeal species, including *Thermococcus kodakarensis* (Briegel et al., 2017), *Pyrococcus furiosus* (Daum et al., 2017), and a closer relative of *H. volcanii*, *Halobacterium salinarum* (Mormile et al., 2003). Although hypothesized to exist (Li et al., 2020), a similar polar structure has not yet been identified in *H. volcanii*, and the components of the polar structures in other archaea are also not known. Together with the observation that CetZ1 is directed to cell poles by MinD proteins (Brown and Duggin, 2024), and strongly localises there in motile cells, our results raise the possibility of a pole-specific or cap-like structure in *H. volcanii* involving CetZ1. CetZ1 appears to polymerize and form protofilaments that can assemble laterally into sheets and are predicted to interact with the inner membrane (de Silva et al., 2024; Duggin et al., 2015). We would predict that such sheet-like structures assemble and help form the structure of the cell poles to promote assembly of motility structures through yet unknown components and interactions.

Both CetZ1 and CetZ2 have even been identified as potential interacting partners with several chemotaxis proteins (including CheW1 and CheY) by affinity purification and mass spectrometry in *H. salinarum* (Schlesner et al., 2012). High resolution structural and compositional analysis of cap structures in *H. volcanii* would be valuable towards identifying any structural role of CetZ1 at cell poles. Since CetZ1, MinD2 (Brown and Duggin, 2024; Patro et al., 2024) and MinD4 (Nußbaum et al., 2020) are all implicated in the positioning of archaella and chemosensory arrays at cell poles, future research could also investigate how these three proteins coordinate their activities to ensure correct positioning and assembly of motility structures.

Overall, our findings suggest that a CetZ family protein participates in multiple distinct roles, and can promote the assembly of other proteins at the cell surface, reminiscent of the roles of tubulin and FtsZ, respectively. Taken together with the diversity and multiplicity of tubulin superfamily proteins in archaea (Aylett and Duggin, 2017; Brown and Duggin, 2023), it is likely that similar characteristics in early archaea led to the evolution of the vast functional complexity, diversity, and specialisation of tubulins in eukaryotes (Brown et al., 2025). Our findings with CetZ1 are also potentially comparable to the fundamental roles of tubulin at the base of the eukaryotic flagellum, which originated before the last eukaryotic common ancestor (Carvalho-Santos et al., 2011; Loreng and Smith, 2017; Mitchell, 2007; Moran et al., 2014). Since flagellar basal body microtubules are derived from mitotic centrioles during eukaryotic cell differentiation, it is interesting to consider parallels to *H. volcanii*, where dynamic repositioning of CetZ1 longitudinal filaments during rod formation (de Silva et al., 2024; Duggin et al., 2015) is followed by the formation of polar structures that might promote the stable assembly of the motility machinery in mature rod cells.

## Experimental Procedures

### Growth and genetic modification of H. volcanii

*H. volcanii* was routinely grown in Hv-Cab or Hv-YPCab medium (de Silva et al., 2021) supplemented with 50 μg/mL uracil if necessary to fulfil auxotrophic requirements of strains, or with L-Tryptophan to induce expression of genes of interest from pTA962-based (Allers et al., 2010) expression plasmids. Cultures were grown at 42 °C with shaking (200 rpm). *H. volcanii* was transformed with demethylated plasmids extracted from *E. coli* strain C2925 using the previously described method (Cline et al., 1989).

*H. volcanii* H26 (ID621) was used as the wildtype background and as the parent strain of the Δ*cetZ1* (ID622) and Δ*cetZ2* (ID357) strains, which were generated as in-frame deletions of *cetZ1* (HVO_2204) or *cetZ2* (HVO_0745) using the previously described plasmids pTA131_2204_IF (Ithurbide et al., 2024) and pTA131_CetZ2 (Duggin et al., 2015), respectively and ‘pop-in pop-out’ method (Allers et al., 2004). Strains ID621, ID622, and ID357 have been confirmed by Illumina whole genome sequencing. For all experiments, *H. volcanii* strains contained the pTA962 shuttle vector (control strains) or vectors for expression of relevant genes. Strains are listed in Table S1.

### Plasmid construction

All reagents for plasmid construction were obtained from New England Biolabs and used according to the manufacturer’s instructions, and oligonucleotides were obtained from Integrated DNA Technologies. Plasmids and oligonucleotides used for cloning detailed in Tables S2 and S3, respectively. To generate a plasmid for the expression of CetZ2-EG-mTurquoise2, the *cetZ2* ORF was amplified from pTA962-CetZ2 using a reverse primer to remove the stop codon and incorporate a BamHI restriction site to the 3’ end. The amplified ORF was then ligated between NdeI and BamHI in pHVID21 (Ithurbide et al., 2024).

For plasmids for dual expression of untagged CetZs and GFP tagged motility proteins, ORFs for ArlD1-GFP, GFP-CheW1, and CheY-GFP were amplified from pSVA3919, pSVA5031, and pSVA5611, respectively, incorporating a BglII restriction site at the 5’ end, and a NotI site at the 3’ end. After purification and restriction digestion, these ORFs were ligated between BamHI and NotI sites of pTA962-CetZ1, pTA962-CetZ2, pTA962-CetZ1.E218A, and pTA962-CetZ2.E212A, utilising the compatible ends generated by BglII and BamHI. The resulting vectors contained the p*.tna* promotor, immediately followed by the CetZ ORF and then the ArlD1, CheW1, or CheY-GFP fusion ORF which was maintained in-frame. The sequences of all constructed plasmids were confirmed by Sanger sequencing (Australian Genome Research Facility).

Vectors for tandem expression of ArlD1-GFP, GFP-CheW1, or CheY-GFP with CetZ1-mCherry (mCh) were generated (pHJB69-71, respectively) as described previously (Ithurbide et al., 2024). The CetZ1-G-mCh fusion (including the flexible Glycine-rich linker) was used here as it complements the motility and shape defects resultant from deletion of *cetZ1* like the CetZ1-G-mTq2 fusion used above (Ithurbide et al., 2024). Briefly, ORFs for ArlD1-GFP, GFP-CheW1, and CheY-GFP were amplified from pSVA3919, pSVA5031, and pSVA5611, respectively, introducing an XbaI restriction site at the 5’ end. These were ligated between NheI and NotI of pHVID132, the vector containing the CetZ1-G-mCh ORF.

### Soft-agar motility assays

To prepare cultures for soft-agar motility assays, single colonies were inoculated into Hv-Cab medium with 1 mM L-Tryptophan and grown to an approximate OD_600_ of 0.6 overnight. Following this, cultures were passaged into fresh Hv-Cab medium with 1 mM L-Tryptophan every 24 hours for three days. All cultures were normalised to the same starting OD_600_ before spotting 2 μL of culture onto the soft agar.

Hv-Cab soft-agar containing 1 mM L-Tryptophan was prepared the day before inoculation of cultures onto the agar. For experiments quantifying halo size, 0.25 % (w/v) agar was used, and motility plates were incubated for 3 days at 42 °C sealed in a plastic bag. This concentration of agar resulted in better uniformity of halo shape and reproducibility of halo size, which was quantified by taking the mean of two perpendicular measurements of the halo diameter. To determine the rate of halo expansion, the radii of motility halos were determined as half the diameter measurements, which taken every 12 hours over 120 hours of total incubation. A linear function was fitted to the data between 60 and 108 hours to determine the slope, representing the rate of halo expansion in mm/h. Motility assays prepared for subsequent microscopy of halo edge cells used 0.3% agar instead, and were incubated for 4-5 days at 42 °C. This method sometimes resulted in irregularly shaped motility halos but produced a more defined leading edge for easier extraction of cells from the agar for analysis by microscopy.

### Light microscopy

Cells were extracted from the edge of motility halos by stabbing the agar with a micropipette (5-10 μL) near the bottom of the agar plate. After pipetting up and down several times to liquify the soft-agar and release the cells, the sample was transferred to a glass slide with a pad containing 1.5 % agarose and 18 % Buffered Salt Water (BSW) (Allers et al., 2004) before covering with a #1.5 glass coverslip. All phase-contrast and fluorescence microscopy were conducted on a V3 DeltaVision Elite inverted microscope with filter sets for GFP/FITC (excitation=464-492 nm, emission=500-523 nm) and mTurquoise2/CFP (excitation=400-453 nm, emission=463-487 nm), an 100X Olympus UPLSAPO/NA 1.4 objective, and pco.edge 5.5 sCMOS camera. Exposure settings were always maintained between all strains expressing the same fusion protein.

### Light microscopy image analysis

To analyse cell shape characteristics and analysis of fluorescence and localization of fluorescent fusion proteins within cells, MicrobeJ v5.13I (Ducret et al., 2016) was used, a plugin for FIJI v2.14.0/1.54f (Schindelin et al., 2012). Unless otherwise mentioned, all settings in MicrobeJ were left as default. Phase-contrast images were used to detect cell outlines using the default thresholding algorithm and a cell area restriction of 1.0-max μm^2^. Cell shape measurements such as area, length, width, aspect ratio, circularity, and longitudinal axis curvature (‘Curvature’ in MicrobeJ) were extracted for individual cells from the MicrobeJ interface. Fluorescent localization was analysed using the *point* detection method. Due to large variance in focus intensity and background fluorescence between genetic backgrounds and plasmids, the tolerance parameter had to be adjusted for each strain as needed for accurate foci detection, but foci detection parameters were maintained between independent replicates. Heatmaps of normalized focus localization were generated using the *XYCellDensity* function, and all other image analysis outputs were extracted from the MicrobeJ results tables.

To conduct local cell curvature analysis and its correlation with fluorescence intensity, a FIJI macro and Python scripts were written and used, based on a previous approach (Duggin et al., 2015). In short, local curvature was analysed using phase-contrast images and the Contour Curvature v1.1.x (www.gluender.de) plugin in FIJI with the number of contour samples (positions) set to 100, and number of coefficient pairs set to 10. The local curvature (1/radius) was recorded at each contour sample position on the outline, and these values were used to calculate the relative standard deviation (RSD) of curvature for each cell. The mean fluorescence intensity along the cell outline generated by Contour Curvature was measured using a 3-pixel line width and was correlated with local curvature by calculating the Pearson correlation coefficient between fluorescence intensity and local curvature for each individual cell. The maximum curvature at each pole was obtained by extracting the largest and second largest local maxima values from curvature vs perimeter position plots. A summary of the image acquisition and analysis pipeline for light microscopy is provided in Fig. S7.

To determine the degree to which polar localised motility markers co-occurred or overlapped with CetZ1, 100 foci of each motility marker were manually identified (blind to CetZ1-mCh), and categorized into one of three patterns with respect to CetZ1-mCh foci: (i) no focus of CetZ1-mCh was located at the same cell pole, (ii) the motility marker and CetZ1-mCh foci were fully or partially overlapping, or (iii) the motility marker and CetZ1-mCh foci were located at the same pole but are not overlapping. This was carried out separately for each independent culture replicate and the mean (and SEM) were obtained.

Using the same fluorescence datasets, MicrobeJ was used to score the occurrence of CetZ1-mCh, ArlD1-GFP, GFP-CheW1, and CheY-GFP in the polar or sub-polar regions. Cell outlines were detected using the phase-contrast channel as described above, and fluorescent foci were detected using the *point* detection method. For the motility markers (GFP channel), tolerance was set to 200, and for CetZ1-mCh tolerance was set to 100. The number of foci per pole were extracted for each channel (For marker-GFP: *Maxima1.count.pole.1* and *Maxima1.count.pole.2*; and for CetZ1-mCh: *Maxima2.count.pole.1* and *Maxima2.count.pole.2*). These were used to determine the number of poles with either one, or both GFP and mCh foci present. We calculated the expected percentage of cell poles with both marker-GFP and CetZ1-mCh foci assuming their co-occurrence is uncorrelated by multiplying the percentage of total poles with at least one GFP focus by the percentage of total poles with at least one mCh focus. The mean, SD and statistical significance were calculated based on the culture replicates.

### Cryo-electron microscopy and analysis

To maximize the number of cells per sample for cryo-electron microscopy, cultures were grown from a theoretical starting OD_600_ of 0.0005 in Hv-YPCab (de Silva et al., 2021) medium and harvested at OD_600_∼0.25 (after ∼24 hours). A 4.5 µL droplet of the culture was applied to glow discharged copper grids (Quantifoil R2/2, Quantifoil Micro Tools) and blotted for 2.5 s in a 95% humidity chamber at 20 °C. The grids were then plunged into liquid ethane using a Lecia EM GP device. The grids were imaged using a Talos Arctica cryo-TEM (Thermo Fisher Scientific), operating at 200 kV with the specimen held at liquid nitrogen temperature. Images were captured at 17,500X magnification using a Falcon 3EC direct electron detector in linear mode. The number of surface filaments per cell (Fig. 1d, e) was manually counted using Cryo-TEM images. This was conducted independently and blindly by three individuals.

## Acknowledgements

The authors would like to acknowledge S. Albers for providing strains and plasmids, as well as L. Cole and A. Bottomley from the UTS Microbial Imaging Facility. We acknowledge the use of the Cryo-Electron Microscopy Facility through the Victor Chang Cardiac Research Institute Innovation Centre, funded by the NSW government and the Electron Microscope Unit within the Mark Wainwright Analytical Centre (MWAC) at UNSW Sydney. The work was supported by the Australian Research Council (DP160101076 and FT160100010 to IGD).

## Author contributions

Conceptualisation – HJB & IGD; Data curation – HJB, MII & JR; Formal analysis – HJB, MII & JR; Investigation – HJB, MII, JR; Methodology – all authors; Supervision – IGD, SI & MABB; Original draft – HJB; Revision – HJB & IGD; Reviewing the manuscript – all authors. Funding Acquisition – IGD.

## Notes

### Competing Interest Statement

The authors have declared no competing interest.

### Summary of Updates

New data included on the measurement of motility over time (overall swimming speed) for wild-type and selected mutants. Extensive further analyses on cell shape, with the addition of further culture replicate datasets. Reorganisation of some results figures and clarification of the main text. New data in the supplementary data figures.

## References

Albers, S.-V., and K.F. Jarrell. 2015. The archaellum: how Archaea swim. Frontiers in Microbiology. 6:23.

Albers, S.-V., and K.F. Jarrell. 2018. The archaellum: an update on the unique archaeal motility structure. Trends in microbiology. 26:351–362.

Allers, T., S. Barak, S. Liddell, K. Wardell, and M. Mevarech. 2010. Improved strains and plasmid vectors for conditional overexpression of His-tagged proteins in Haloferax volcanii. Applied and environmental microbiology. 76:1759–1769.

Allers, T., H.-P. Ngo, M. Mevarech, and R.G. Lloyd. 2004. Development of additional selectable markers for the halophilic archaeon Haloferax volcanii based on the leuB and trpA genes. Applied and environmental microbiology. 70:943–953.

An, S., Y. Deng, J.W. Tomsho, M. Kyoung, and S.J. Benkovic. 2010. Microtubule-assisted mechanism for functional metabolic macromolecular complex formation. Proceedings of the National Academy of Sciences. 107:12872–12876.

Appert-Rolland, C., M. Ebbinghaus, and L. Santen. 2015. Intracellular transport driven by cytoskeletal motors: General mechanisms and defects. Physics Reports. 593:1–59.

Aylett, C.H., and I.G. Duggin. 2017. The tubulin superfamily in archaea. *In* Prokaryotic Cytoskeletons. Springer. 393–417.

Barton, N.R., and L. Goldstein. 1996. Going mobile: microtubule motors and chromosome segregation. Proceedings of the National Academy of Sciences. 93:1735–1742.

Beeby, M., J.L. Ferreira, P. Tripp, S.-V. Albers, and D.R. Mitchell. 2020. Propulsive nanomachines: the convergent evolution of archaella, flagella and cilia. FEMS microbiology reviews. 44:253–304.

Briegel, A., C.M. Oikonomou, Y.W. Chang, A. Kjær, A.N. Huang, K.W. Kim, D. Ghosal, H.H. Nguyen, D. Kenny, and R.R. Ogorzalek Loo. 2017. Morphology of the archaellar motor and associated cytoplasmic cone in Thermococcus kodakaraensis. EMBO reports. 18:1660–1670.

Brown, H.J., and I.G. Duggin. 2023. Diversity and Potential Multifunctionality of Archaeal CetZ Tubulin-like Cytoskeletal Proteins. Biomolecules. 13:134.

Brown, H.J., and I.G. Duggin. 2024. MinD proteins regulate CetZ1 localization in Haloferax volcanii. Frontiers in Microbiology. 15:1474697.

Brown, H.J., V.D. Shinde, L. Bosi, and I.G. Duggin. 2025. Evolution of the cytoskeleton: Emerging clues from the diversification and specialisation of archaeal cytoskeletal proteins. Current opinion in cell biology. 95:102557.

Carvalho-Santos, Z., J. Azimzadeh, J.B. Pereira-Leal, and M. Bettencourt-Dias. 2011. Evolution: Tracing the origins of centrioles, cilia, and flagella. J Cell Biol. 194:165–175.

Cline, S.W., W.L. Lam, R.L. Charlebois, L.C. Schalkwyk, and W.F. Doolittle. 1989. Transformation methods for halophilic archaebacteria. Canadian Journal of Microbiology. 35:148–152.

Daum, B., J. Vonck, A. Bellack, P. Chaudhury, R. Reichelt, S.-V. Albers, R. Rachel, and W. Kühlbrandt. 2017. Structure and in situ organisation of the Pyrococcus furiosus archaellum machinery. Elife. 6:e27470.

de Silva, R.T., M.F. Abdul-Halim, D.A. Pittrich, H.J. Brown, M. Pohlschroder, and I.G. Duggin. 2021. Improved growth and morphological plasticity of Haloferax volcanii. Microbiology.

de Silva, R.T., V. Shinde, H.J. Brown, Y. Liao, and I.G. Duggin. 2024. Dynamic self-association of archaeal tubulin-like protein CetZ1 drives Haloferax volcanii morphogenesis. bioRxiv:2024.2004. 2008.588506.

Ducret, A., E.M. Quardokus, and Y.V. Brun. 2016. MicrobeJ, a tool for high throughput bacterial cell detection and quantitative analysis. Nature microbiology. 1:1–7.

Duggin, I.G., C.H. Aylett, J.C. Walsh, K.A. Michie, Q. Wang, L. Turnbull, E.M. Dawson, E.J. Harry, C.B. Whitchurch, and L.A. Amos. 2015. CetZ tubulin-like proteins control archaeal cell shape. Nature. 519:362.

Erickson, H.P. 1995. FtsZ, a prokaryotic homolog of tubulin? Cell. 80:367–370.

Erickson, H.P. 2007. Evolution of the cytoskeleton. Bioessays. 29:668–677.

Hawkins, T., M. Mirigian, M.S. Yasar, and J.L. Ross. 2010. Mechanics of microtubules. Journal of biomechanics. 43:23–30.

Hirokawa, N., Y. Noda, Y. Tanaka, and S. Niwa. 2009. Kinesin superfamily motor proteins and intracellular transport. Nature reviews Molecular cell biology. 10:682–696.

Ithurbide, S., R.T. de Silva, H.J. Brown, V. Shinde, and I.G. Duggin. 2024. A vector system for single and tandem expression of cloned genes and multi-colour fluorescent tagging in Haloferax volcanii. Microbiology (Reading). 170:001461.

Jarrell, K.F., and S.-V. Albers. 2012. The archaellum: an old motility structure with a new name. Trends in microbiology. 20:307–312.

Jarrell, K.F., S.-V. Albers, and J. Machado. 2021. A comprehensive history of motility and Archaellation in Archaea. FEMS Microbes. 2.

Kupper, J., W. Marwan, D. Typke, H. Grünberg, U. Uwer, M. Gluch, and D. Oesterhelt. 1994. The flagellar bundle of Halobacterium salinarium is inserted into a distinct polar cap structure. Journal of bacteriology. 176:5184–5187.

Li, Z., Y. Kinosita, M. Rodriguez-Franco, P. Nußbaum, F. Braun, F. Delpech, T.E. Quax, and S.-V. Albers. 2019. Positioning of the motility machinery in halophilic archaea. MBio. 10:e00377–00319.

Li, Z., M. Rodriguez-Franco, S.V. Albers, and T.E. Quax. 2020. The switch complex ArlCDE connects the chemotaxis system and the archaellum. Molecular microbiology. 114:468–479.

Loreng, T.D., and E.F. Smith. 2017. The Central Apparatus of Cilia and Eukaryotic Flagella. Cold Spring Harb Perspect Biol. 9.

Lutkenhaus, J., S. Pichoff, and S. Du. 2012. Bacterial cytokinesis: from Z ring to divisome. Cytoskeleton. 69:778–790.

Mitchell, D.R. 2007. The evolution of eukaryotic cilia and flagella as motile and sensory organelles. Adv Exp Med Biol. 607:130–140.

Moran, J., P.G. McKean, and M.L. Ginger. 2014. Eukaryotic Flagella: Variations in Form, Function, and Composition during Evolution. BioScience. 64:1103–1114.

Mormile, M.R., M.A. Biesen, M.C. Gutierrez, A. Ventosa, J.B. Pavlovich, T.C. Onstott, and J.K. Fredrickson. 2003. Isolation of Halobacterium salinarum retrieved directly from halite brine inclusions. Environmental Microbiology. 5:1094–1102.

Nogales, E., K.H. Downing, L.A. Amos, and J. Löwe. 1998. Tubulin and FtsZ form a distinct family of GTPases. Nature Structural and Molecular Biology. 5:451.

Nußbaum, P., S. Ithurbide, J.C. Walsh, M. Patro, F. Delpech, M. Rodriguez-Franco, P.M. Curmi, I.G. Duggin, T.E. Quax, and S.-V. Albers. 2020. An oscillating MinD protein determines the cellular positioning of the motility machinery in archaea. Current Biology. 30:4956–4972. e4954.

Patro, M., F. Grünberger, S. Sivabalasarma, S. Gfrerer, M. Rodriguez-Franco, P. Nußbaum, D. Grohmann, S. Ithurbide, and S.-V. Albers. 2024. MinD2 modulates cell shape and motility in the archaeon Haloferax volcanii. Frontiers in Microbiology. 15:1474570.

Prelich, G. 2012. Gene overexpression: uses, mechanisms, and interpretation. Genetics. 190:841–854.

Quax, T.E., F. Altegoer, F. Rossi, Z. Li, M. Rodriguez-Franco, F. Kraus, G. Bange, and S.-V. Albers. 2018. Structure and function of the archaeal response regulator CheY. Proceedings of the National Academy of Sciences. 115:E1259–E1268.

Schindelin, J., I. Arganda-Carreras, E. Frise, V. Kaynig, M. Longair, T. Pietzsch, S. Preibisch, C. Rueden, S. Saalfeld, and B. Schmid. 2012. Fiji: an open-source platform for biological-image analysis. Nature methods. 9:676.

Schlesner, M., A. Miller, H. Besir, M. Aivaliotis, J. Streif, B. Scheffer, F. Siedler, and D. Oesterhelt. 2012. The protein interaction network of a taxis signal transduction system in a halophilic archaeon. BMC microbiology. 12:1–20.

Schwarzer, S., M. Rodriguez-Franco, H.M. Oksanen, and T.E. Quax. 2021. Growth phase dependent cell shape of Haloarcula. Microorganisms. 9:231.

Vaughan, S., B. Wickstead, K. Gull, and S.G. Addinall. 2004. Molecular evolution of FtsZ protein sequences encoded within the genomes of archaea, bacteria, and eukaryota. Journal of molecular evolution. 58:19–29.

Young, K.D. 2006. The selective value of bacterial shape. Microbiology and molecular biology reviews. 70:660–703.

