## Supplementary Information for "CetZ1-dependent polar assembly of the archaeal motility machinery"

### Contents

**Supplementary Methods**

**Supplementary Figures**

1. Motility of various *cetZ* knockout and overexpression strains.
2. Cell morphology phenotypes of *cetZ* mutants.
3. Dose-dependent effects of *cetZ1* and *cetZ1.E218A* overexpression on motility.
4. Surface adhesion of *cetZ1/2* deletion and overexpression strains.
5. Tryptophan-induced expression levels of CetZs and CetZ-mTq2 fusions.
6. Motility and cell shape assays of various motility marker deletion and GFP fusion expression strains.
7. Extended localisation analysis of ArlD1-GFP, GFP-CheW1, and CheY-GFP.
8. Workflow to image and analyse foci localisation and cell curvature.
9. Extended cell shape analysis of ArlD1-GFP, GFP-CheW1, and CheY-GFP expressing strains.

**Supplementary Tables**

1. Strains used in this study
2. Plasmids used in this study
3. Oligonucleotides used in this study

**References**

**Supplementary Methods**

***Growth curves***

Growth curves were generated by starting two replicate cultures of various strains started from single colonies. Cells were grown in Hv-Cab medium supplemented with 1 mM L-tryptophan and passaged twice before equalising all cultures to $OD_{600}$~0.01 in fresh Hv-Cab medium with 1 mM L-tryptophan. Each culture was aliquoted twice, generating technical duplicates. Growth of these cultures was carried out in a 96-well plate using the Tecan Spark 20M plate reader, incubating at 42 °C with shaking at 500 rpm, taking absorbance readings at 600 nm every 30 minutes.

***Surface adhesion assay***

Surface adhesion was measured using a previously described method [1]. Three culture replicates of each strain were grown overnight in Hv-Cab or Hv-YPCab medium supplemented with 1 mM L-Tryptophan. Culture replicates were then diluted to a theoretical starting $OD_{600}$ of 0.3 in fresh medium before aliquoting into a 96-well flat-bottom plate in triplicate. Using a Tecan Spark® 20M microplate reader/incubator, cultures were incubated statically at 42°C for a total of 32 hours, introducing shaking at 200 rpm for a duration of 4 minutes every 20 minutes. Following this, the culture medium was carefully discarded, and cells were fixed using 2% (v/v) acetic acid for 3 minutes. Fixed cells were air-dried and stained with 0.1% (w/v) crystal violet for 10 min, followed by three washes with distilled water. Finally, 10% (v/v) acetic acid and 30% (v/v) methanol was used to release the crystal violet stain, and absorbance at $OD_{600}$ of each well was measured in the microplate reader.

***SDS-PAGE and western blotting***

Samples for SDS-PAGE were prepared by resuspending whole-cell pellets in a lysis buffer (20 mM Tris-HCl pH 7.5), 1X DNAse 1, 1X EDTA-free protease inhibitor). After incubating for 30 min at room temperature, 4X SDS-PAGE sample buffer (250 mM Tris-HCl pH 6.8, 40% (v/v) glycerol, 4% (w/v) SDS, 0.04% (w/v) Bromophenol blue, 20% (v/v) 2-mercaptoethanol) was added, bringing samples to a theoretical $OD_{600}$ of 5. Samples were vortexed thoroughly before loading onto a 4-20% Mini-PROTEAN® TGX™ pre-cast polyacrylamide gel (Bio-Rad), using the Broad Range Blue Prestained Protein Standard (New England Biolabs). Electrophoresis was carried out at 100 V for 85 min with a Bio-Rad Mini-Protean3 electrophoresis system.

Protein was transferred to a nitrocellulose membrane using the Trans-Blot Turbo mini transfer system (Bio-Rad), followed by total protein staining with Ponceau S (0.1% (w/v) in 5% (v/v) acetic acid). Blocking of the membrane was carried out using 5% (w/v) skim milk powder in TBST (50 mM Tris-HCl, pH 7.4, 150 mM NaCl, 0.05% (v/v) Tween-20) for 3 h at room temperature. Then, incubation with primary antibodies against CetZ1 or CetZ2, diluted 1/1000 in TBST, was done overnight at 4°C. Following this, the membrane was washed using TBST, and incubated with the secondary antibody (donkey anti-rabbit IgG HRP conjugate, Abcam 16284) for 1 h at room temperature, used as a 1/5000 dilution in TBST. After washing, the membrane was incubated with the SuperSignal™ West Pico PLUS chemiluminescent substrate (ThermoFisher) for 5 min and imaged using the Amersham™ Imager 600 (GE Healthcare).

### Supplementary Figures

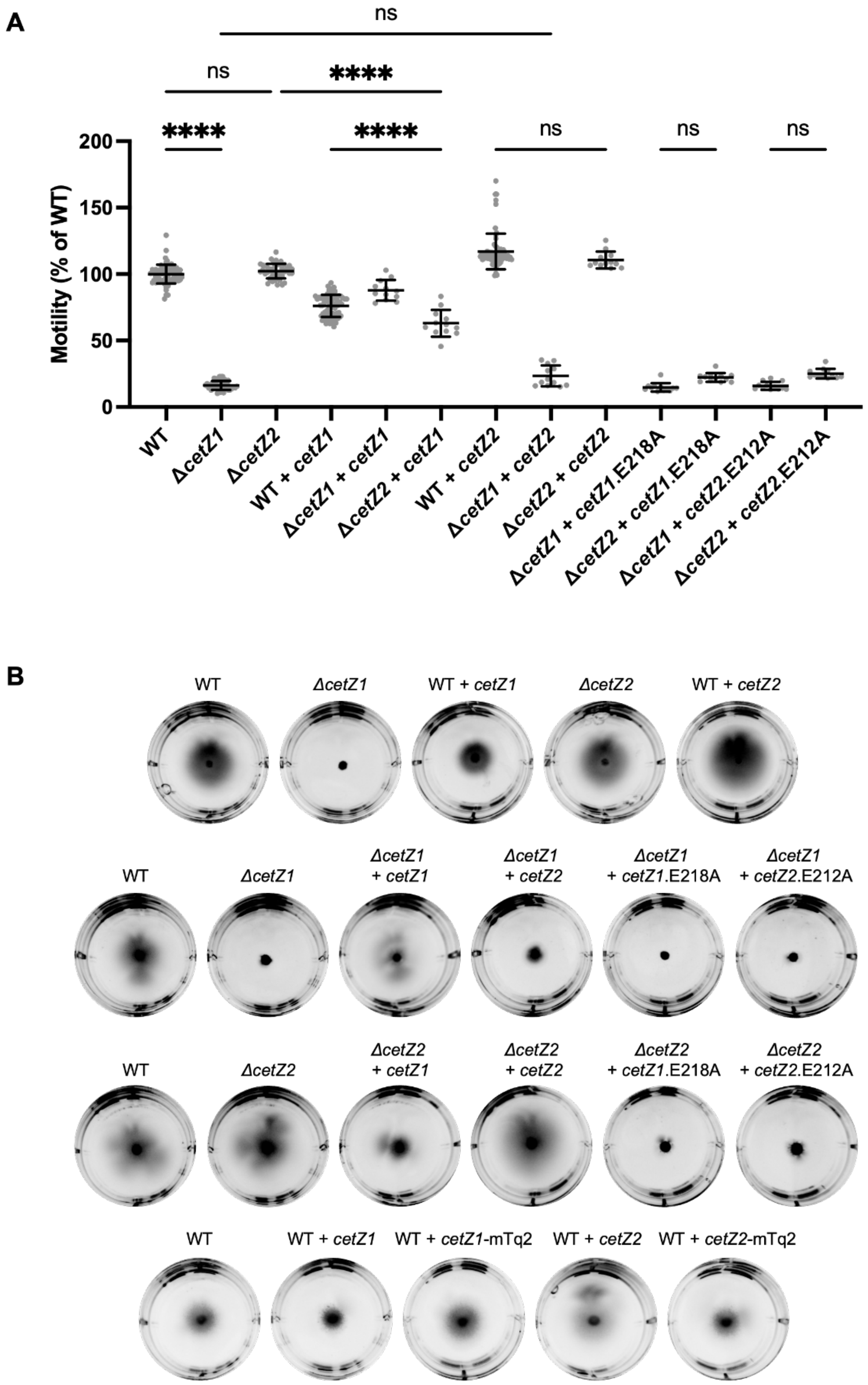


**Figure S1. Motility of various *cetZ* knockout and overexpression strains. A)** Quantification of motility halo diameter from Hv-Cab soft-agar (0.25%) motility assays with 1 mM L-Tryptophan. Individual points represent biological replicates, mean and standard deviation shown. One-way ANOVA was used as a statistical test, where ns = not significant, ****p<0.0001, and only relevant comparisons are shown. **B)** Representative examples of motility halos of strains in **A.** Data from Figure 1a and b is included in panels **A** and **B**, respectively, for reference.

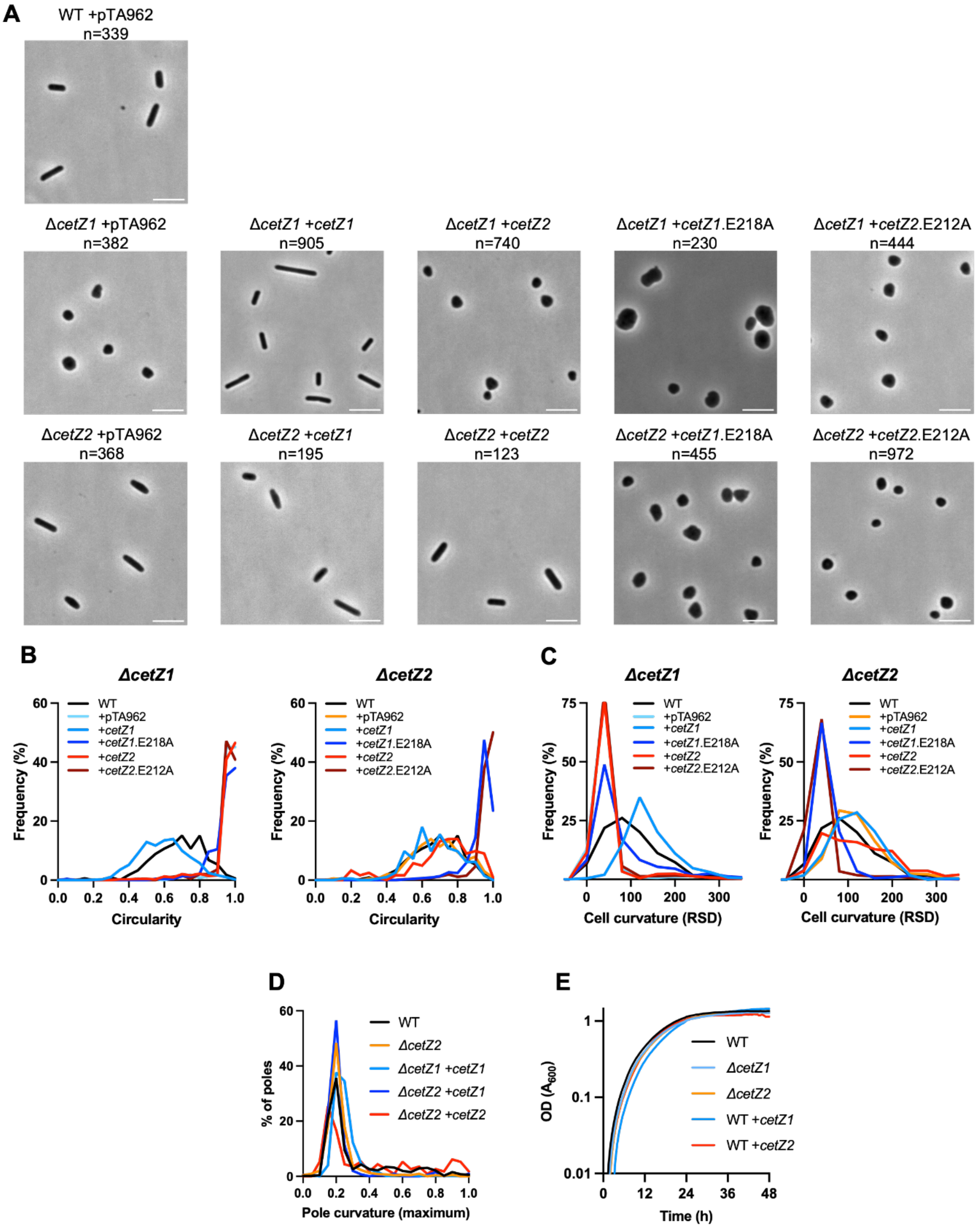


**Figure S2. Cell elongation (circularity) during expression of various *cetZ1/2* or their indicated mutants in *cetZ* knock-out backgrounds. A)** Phase-contrast microscopy of cells extracted from the halo edge. Scale bar: 5 µm. These images were used to quantify **B)** cell circularity, represented as a frequency distribution, and **C)** relative standard deviation (RSD) of curvature of the cell outlines of individual cells. In panels **B** and **C,** data for WT (H26 with pTA962) is shown on all graphs for reference. **D)** Frequency distribution showing maximum local curvature of cell poles for rod-forming strains. The number of individual cells imaged (**A**), and analysed (**B-D**) is indicated in panel **A**. **E)** Equivalent growth rates (see supplementary methods) of key *cetZ* mutants, in Hv-Cab medium supplemented with 1 mM L-Tryptophan.

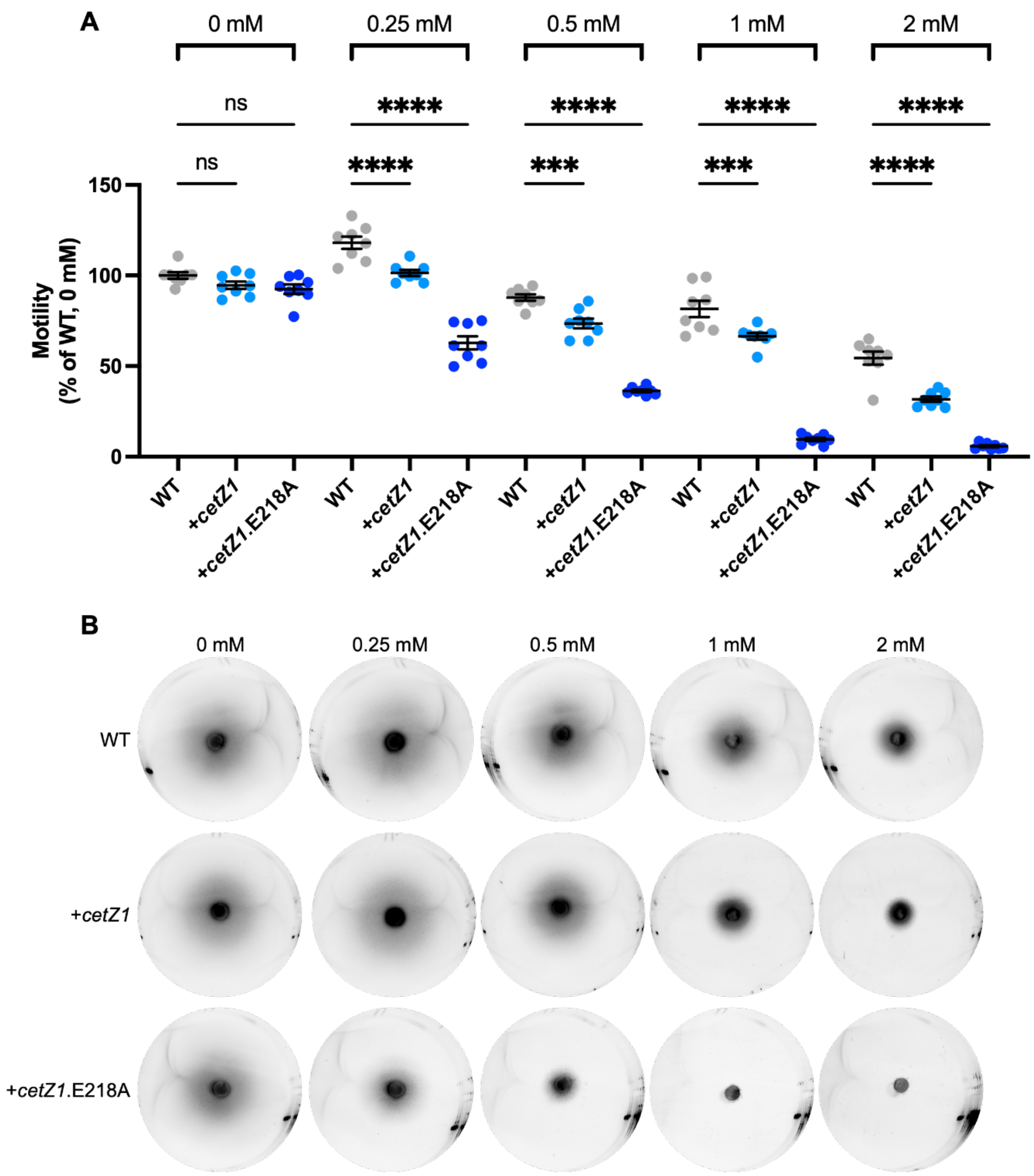


**Figure S3. Dose-dependent effects of *cetZ1* and *cetZ1.E218A* overexpression on motility.** From pTA962-*cetZ1* and pTA962-*cetZ1*.E218A, *cetZ1* and *cetZ1*.E218A were expressed in H26 wildtype, using varying concentrations (0, 0.25, 0.5, 1, or 2 mM) of L-Tryptophan to induce expression. **A)** Quantification of motility halo diameters. Individual points represent biological replicates, mean and standard deviation are shown. One-way ANOVA was used as a statistical test, ****p<0.0001, ***p<0.0002, ns=not significant. **B)** Representative motility halos for strains and L-Tryptophan concentrations in **A**.

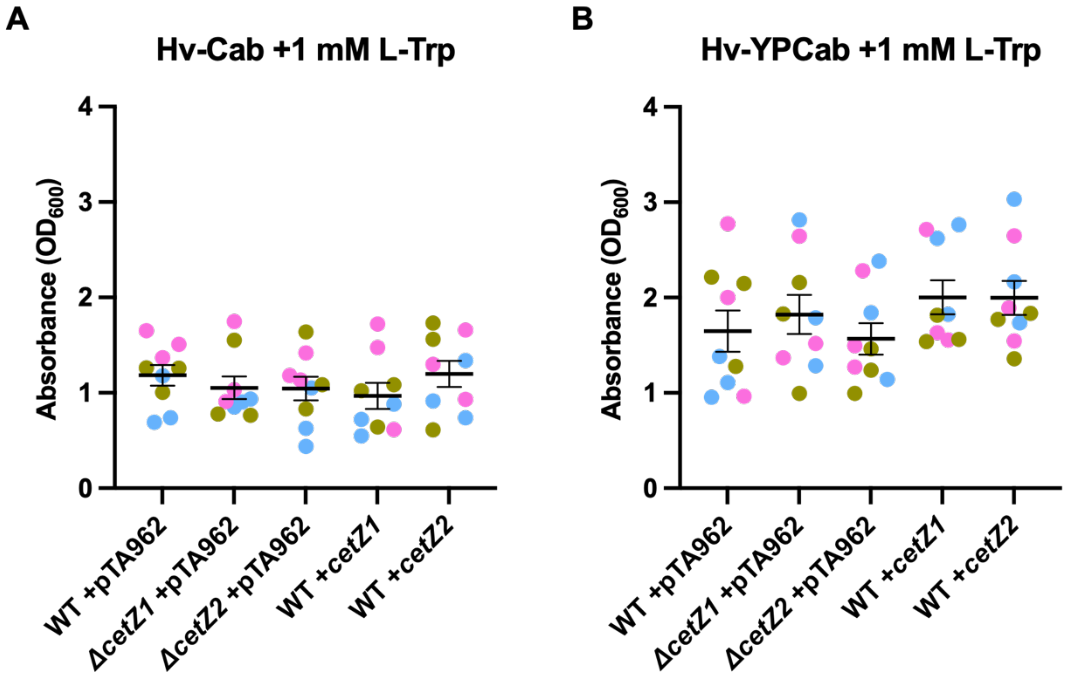


**Figure S4. Surface adhesion of *cetZ1/2* deletion and overexpression strains.** The absorbance at 600 nm was measured after crystal violet straining of surface bound cells (see supplementary methods). Three culture replicates were conducted for each strain (coloured magenta, cyan, and yellow), and each culture replicate was carried out in triplicate, in both **A)** Hv-Cab and **B)** Hv-YPCab medium supplemented with 1 mM L-Tryptophan. One-way ANOVA was used as a statistical test, comparing *cetZ1/2* deletion and overexpression strains to H26 wildtype, however all comparisons were not statistically significant.

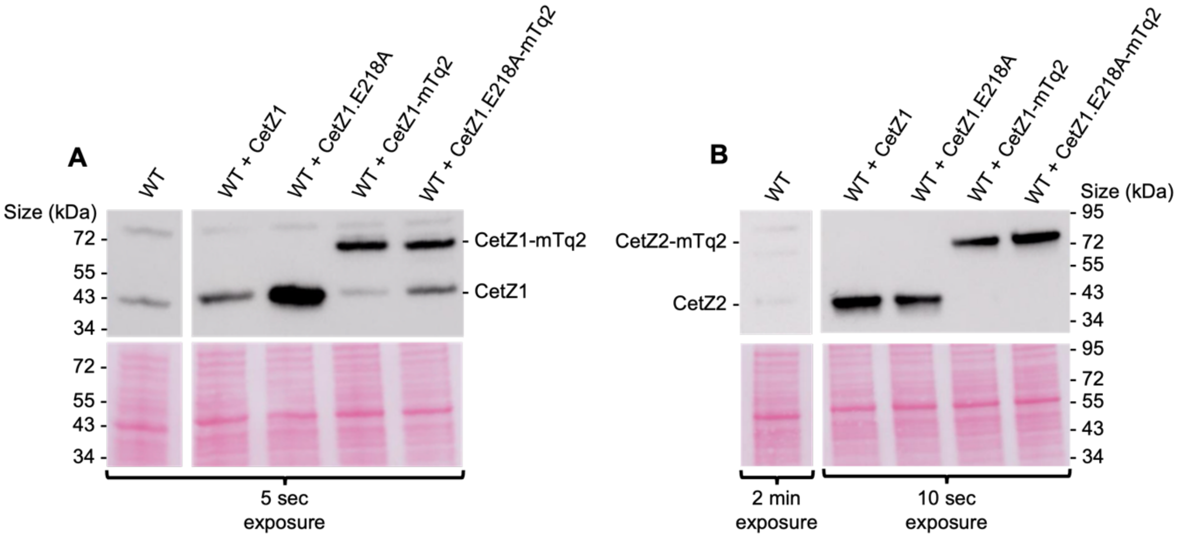


**Figure S5. Tryptophan-induced expression levels of CetZs and CetZ-mTq2 fusions.** Western blotting using cells growing in batch culture, sampled during mid-log phase. Whole cell lysates were loaded onto an SDS-PAGE gel for subsequent western blotting (see supplementary methods). Western blots using **A)** anti-CetZ1 and **B)** anti-CetZ2 antibody. Lower panels show total protein (Ponceau S) staining.

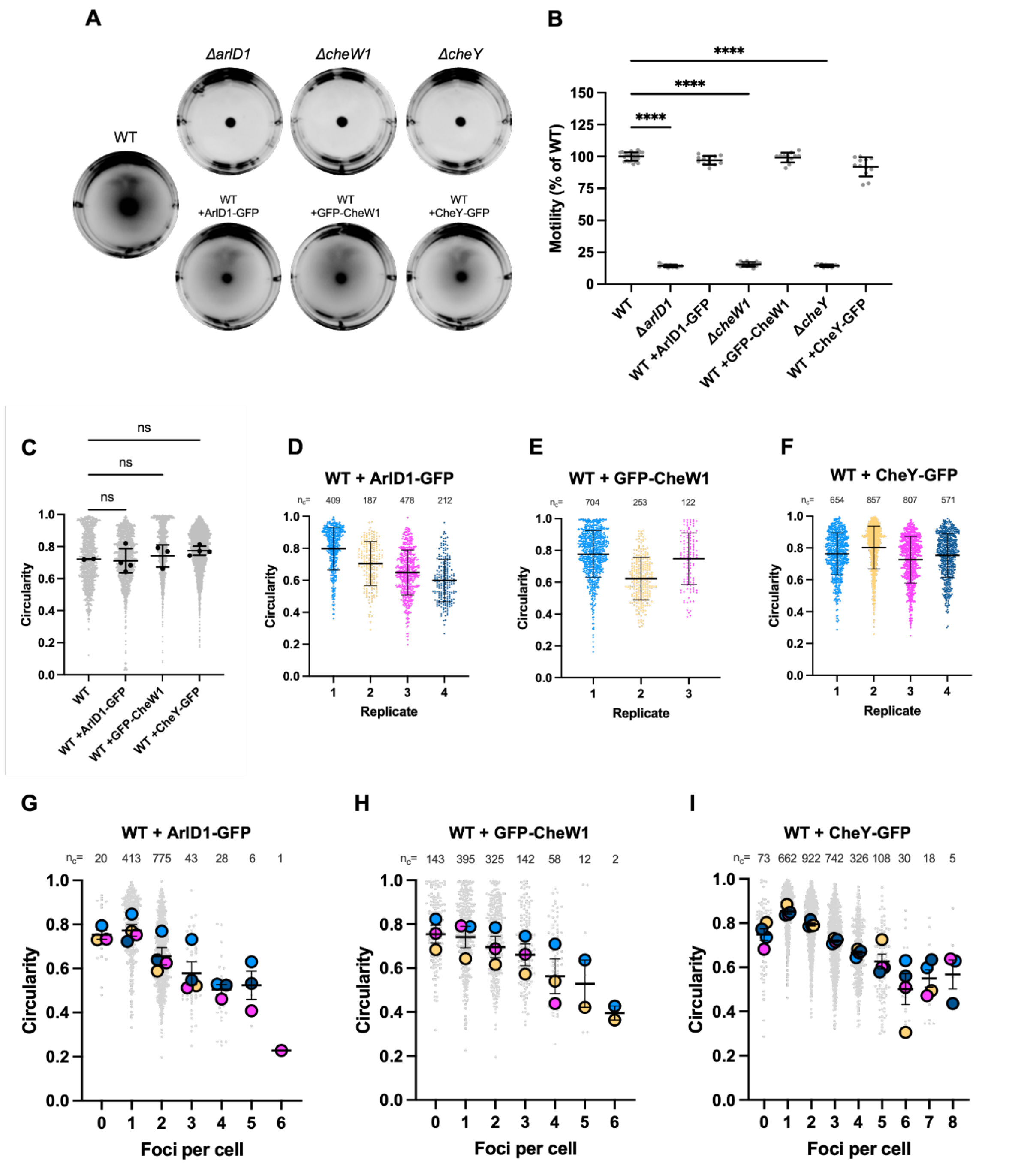


**Figure S6. Motility and cell shape assays of various motility marker deletion and GFP fusion expression strains. A)** Representative images and **B)** quantification of motility halo diameter of H26 wildtype cells or *arlD1*, *cheW1*, and *cheY* deletion strains with pTA962 or H26 wildtype cells expressing ArlD1, CheW1, or CheY fusions with GFP. Individual points represent independent replicates. **C)** Cell circularity of cells withdrawn from the leading edge of motility halos and imaged with phase-contrast microscopy (shown in Fig. 5a, 6a, and 7a, respectively). Individual points represent the mean circularity of all cells within one biological replicate. The number of cells measured from pooled biological replicates is as follows: WT +pTA962, n=1078; WT +ArlD1-GFP, n=1269; WT +GFP-CheW1, n=1093 WT + CheY-GFP, n=2950. **D-F)** Cell circularity displayed individually for each biological (culture) replicate. Individual points represent individual cells, and points are coloured by replicate and the number of individual cells in each replicate are indicated above the data. **G-I)** Circularity by number of foci detected. Large, coloured data points indicate mean cell circularity for individual replicates as in **D-F**. Small grey data points indicate individual cells, pooled from all biological replicates, and the number of individual cells in each category are indicated above the data. In **B** and **C-I**, mean and standard deviation is shown. One-way ANOVA was used as a statistical test. ****p<0.0001, ns=not significant. Motility data from Figures 4a, 6a, and 7a is included in panel **A** for reference. Circularity data from Figures 5c, 6c, and 7c is included in panel **C,** and used to generate panels **D-I.**

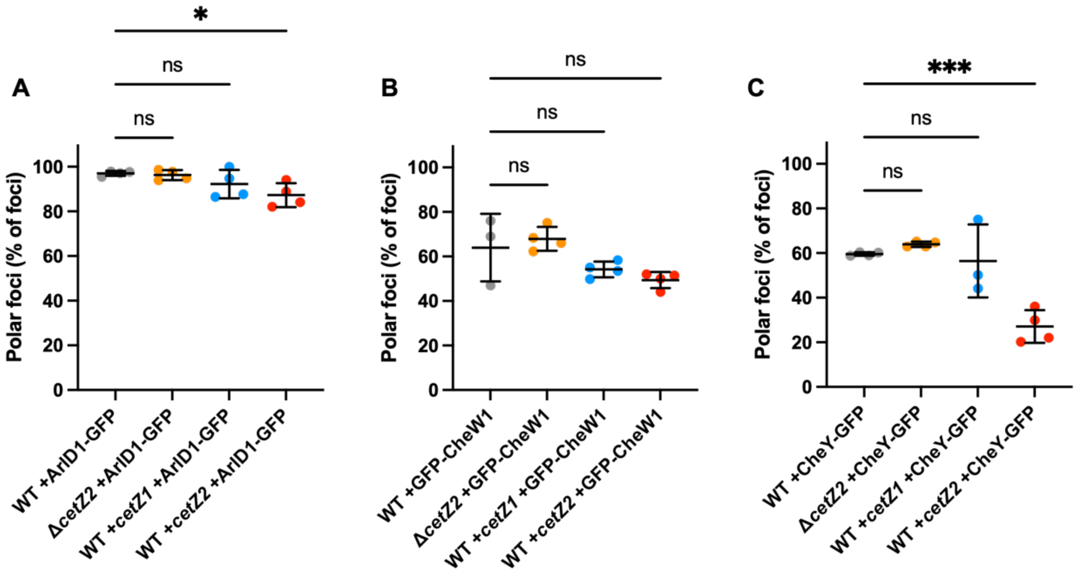


**Figure S7. Extended localisation analysis of ArlD1-GFP, GFP-CheW1, and CheY-GFP.** Proportion of detected foci located at cell poles for ArlD1-GFP, GFP-CheW1, and CheY-GFP, respectively. In all panels, individual points represent the mean of one biological (culture) replicate. Error bars indicate mean and standard deviation. One-way ANOVA was used as a statistical test. ***p<0.0002, ns=not significant.

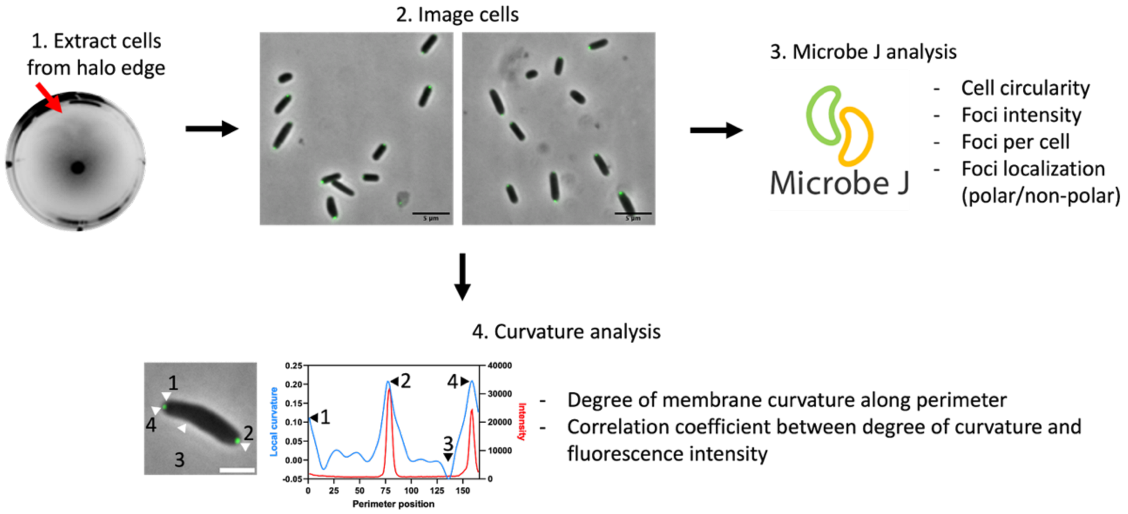


**Figure S8. Workflow to image and analyse foci localisation and cell curvature.** (1) First, cultures were inoculated onto Hv-Cab soft-agar (0.3%) supplemented with 1 mM L-Tryptophan and allowed to grow at 42°C. After 4-5 days (once a clearly defined leading edge was developed by actively swimming cells), cells were extracted from the leading edge of the motility halo and directly spotted onto an agarose pad for (2) analysis by phase-contrast and fluorescence microscopy. The examples shown are WT + ArlD1-GFP. (3) The images were then analysed in MicrobeJ as described in the methods section. Phase-contrast images were used to detect cell outlines and measure cell circularity, while the fluorescence microscopy images were used for foci analysis. Focus intensity counts of foci per cell, foci localisation at cell poles, and heat map representations of foci localisation were all obtained from the MicrobeJ results interface. (4) The same images obtained in step 2 were subjected to curvature and intensity correlation analysis, where plots of local curvature (obtained from phase-contrast images), and fluorescence intensity (obtained from fluorescence images) along the cell perimeter were generated for individual cells, and the Pearson's Correlation Coefficient measured. Local curvature plots (4, blue line) were also used to identify maximum pole curvature (Fig. S9b, e, h).

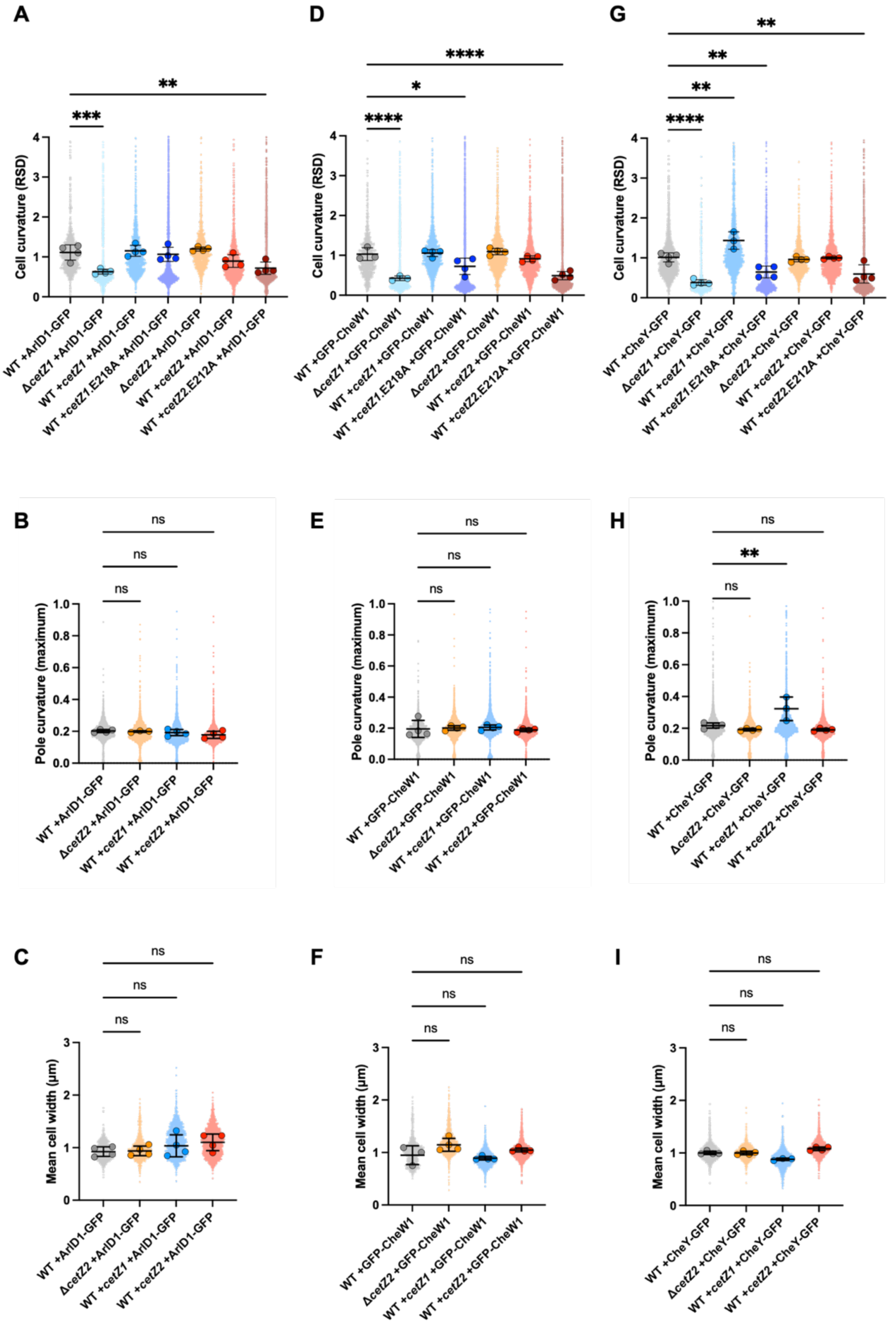


**Figure S9. Extended cell shape analysis of ArlD1-GFP, GFP-CheW1, and CheY-GFP expressing strains.** **A, D, G)** Quantification of the relative standard deviation of curvature of the cell outlines for various *cetZ*

mutants expressing ArlD1-GFP, GFP-CheW1, or CheY-GFP. This measurement represents overall variance in curvature of each cell's outline, and was calculated like described previously [2]. **B, E, H)** Quantification of the maximum curvature at cell poles in rod-forming strains. **C, F, I)** Quantification of mean cell width for rod-forming *cetZ* mutants expressing ArlD1-GFP, GFP-CheW1, and CheY-GFP. All panels are represented as superplots, where small individual points represent individual cells or poles, and large data points represent the mean for one culture replicate. Error bars indicate mean and standard deviation of culture replicates. One-way ANOVA was used as a statistical test. ****p<0.0001, ***p<0.0002, **p<0.002, *p<0.02, ns=not significant.

### Supplementary Tables

**Table S1.** Strains used in this study

| Strain | Description | Source |
|---|---|---|
| ID621 (H26) | Wildtype strain, cured of pHV2 and *ΔpyrE2* | [3] |
| ID622 (Δ*cetZ1*) | H26 with deletion of *cetZ1* (HVO_2204) | This study |
| ID357 (Δ*cetZ2*) | H26 with deletion of *cetZ2* (HVO_0745) | This study |
| Δ*arlD1* | H26 with deletion of *arlD1* (HVO_1203) | [4] |
| Δ*cheW1* | H26 with deletion of *cheW1* (HVO_1225) | [4] |
| Δ*cheY* | H26 with deletion of *cheY* (HVO_1207) | [5] |

**Table S2.** Plasmids used in this study

| Plasmid | Description | Source |
|---|---|---|
| pTA131_2204_IF | pTA131[3] containing flanking sequences for in-frame deletion of *cetZ1* (HVO_2204). | [6] |
| pTA131_CetZ2 | pTA131[3] containing flanking sequences for in-frame deletion of *cetZ2* (HVO_0745). | [2] |
| pTA962 | Expression vector containing *p.tna* promotor for *H. volcanii* and *E. coli* shuttle plasmid. | [7] |
| pHVID21 | For construction of C-terminal EG-linker mTurquoise2 fusion proteins. | [6] |
| pTA962-CetZ1 | For expression of CetZ1 under the control of the *p.tna* promotor. | [2] |
| pTA962-CetZ2 | For expression of CetZ2 under the control of the *p.tna* promotor. | [2] |
| pTA962-CetZ1.E218A | For expression of CetZ1.E218A under the control of the *p.tna* promotor. | [2] |
| pTA962-CetZ2.E212A | For expression of CetZ2.E212A under the control of the *p.tna* promotor. | [2] |
| pHVID135 | For expression of CetZ1-G-mTurquoise2 C-terminal fusion under the control of the *p.tna* promotor. | [6] |
| pHJB6 | For expression of CetZ2-EG-mTurquoise2 C-terminal fusion under the control of the *p.tna* promotor. | This study |
| pSVA3919 | For expression of ArlD1-GFP C-terminal fusion under the control of the *p.tna* promotor. | [4] |
| pSVA5031 | For expression of GFP-CheW1 N-terminal fusion under the control of the *p.tna* promotor. | [4] |
| pSVA5611 | For expression of CheY-GFP C-terminal fusion under the control of the *p.tna* promotor. | [4] |
| pHJB31 | pTA962-CetZ1 with ArlD1-GFP cloned between BamHI and NotI restriction sites. For dual expression of CetZ1 and ArlD1-GFP under the control of the same *p.tna* promotor. | This study |
| pHJB32 | pTA962-CetZ1 with GFP-CheW1 cloned between BamHI and NotI restriction sites. For dual expression of CetZ1 and GFP-CheW1 under the control of the same *p.tna* promotor. | This study |
| pHJB33 | pTA962-CetZ1 with CheY-GFP cloned between BamHI and NotI restriction sites. For dual expression of CetZ1 and CheY-GFP under the control of the same *p.tna* promotor. | This study |
| pHJB34 | pTA962-CetZ1.E218A with ArlD1-GFP cloned between BamHI and NotI restriction sites. For dual expression of CetZ1.E218A and ArlD1-GFP under the control of the same *p.tna* promotor. | This study |
| pHJB35 | pTA962-CetZ1.E218A with GFP-CheW1 cloned between BamHI and NotI restriction sites. For dual expression of CetZ1.E218A and GFP-CheW1 under the control of the same *p.tna* promotor. | This study |
| pHJB36 | pTA962-CetZ1.E218A with CheY-GFP cloned between BamHI and NotI restriction sites. For dual expression of CetZ1.E218A and CheY-GFP under the control of the same *p.tna* promotor. | This study |

| | | |
|---|---|---|
| pHJB37 | pTA962-CetZ2 with ArlD1-GFP cloned between BamHI and NotI restriction sites. For dual expression of CetZ2 and ArlD1-GFP under the control of the same *p.tna* promotor. | This study |
| pHJB38 | pTA962-CetZ2 with GFP-CheW1 cloned between BamHI and NotI restriction sites. For dual expression of CetZ2 and GFP-CheW1 under the control of the same *p.tna* promotor. | This study |
| pHJB39 | pTA962-CetZ2 with CheY-GFP cloned between BamHI and NotI restriction sites. For dual expression of CetZ2 and CheY-GFP under the control of the same *p.tna* promotor. | This study |
| pHJB40 | pTA962-CetZ2.E212A with ArlD1-GFP cloned between BamHI and NotI restriction sites. For dual expression of CetZ2.E212A and ArlD1-GFP under the control of the same *p.tna* promotor. | This study |
| pHJB41 | pTA962-CetZ2.E212A with GFP-CheW1 cloned between BamHI and NotI restriction sites. For dual expression of CetZ2.E212A and GFP-CheW1 under the control of the same *p.tna* promotor. | This study |
| pHJB42 | pTA962-CetZ2.E212A with CheY-GFP cloned between BamHI and NotI restriction sites. For dual expression of CetZ2.E212A and CheY-GFP under the control of the same *p.tna* promotor. | This study |
| pHVID132 | For expression of CetZ1-G-mCherry C-terminal fusion under the control of the *p.tna* promotor. | [6] |
| pHJB69 | pHVID132 with ArlD1-GFP cloned between NheI and NotI restriction sites. For dual expression of CetZ1-mCh and ArlD-GFP under the control of the same *p.tna* promotor. | This study |
| pHJB70 | pHVID132 with GFP-CheW1 cloned between NheI and NotI restriction sites. For dual expression of CetZ1-mCh and GFP-CheW1 under the control of the same *p.tna* promotor. | This study |
| pHJB71 | pHVID132 with CheY-GFP cloned between NheI and NotI restriction sites. For dual expression of CetZ1-mCh and CheY-GFP under the control of the same *p.tna* promotor. | This study |

**Table S3.** Oligonucleotides used in this study

| Name | Sequence (5'-3') | Use |
|---|---|---|
| cetZ2_RnoS | CGCGGATCCCAGCAGGTCGTCGAGGTCGTCGCT | Reverse primer to amplify the CetZ2 ORF excluding its stop codon, incorporating a BamHI restriction site. |
| BglII_ArlD1 | GGCGGCAGATCTATGGCAAGCAAGGT | Forward primer to amplify the ArlD1-GFP ORF, incorporating BglII restriction site. |
| BglII_GFP | GGCGGCAGATCTATGAGTAAAGGAGAAG | Forward primer to amplify the GFP-CheW1 ORF, incorporating BglII restriction site. |
| BglII_CheY | GGCGGCAGATCTATGGACTCTATCGTCGGCGAC | Forward primer to amplify the CheY-GFP ORF, incorporating BglII restriction site. |
| T7 | TAATACGACTCACTATAGGG | Forward primer used to amplify the CetZ2 ORF from pTA962-CetZ2 for cloning into pHVID21. |
| T3 | GCAATTAACCCTCACTAAAGG | Reverse primer used to amplify the ArlD1-GFP, GFP-CheW1, and CheY-GFP ORFs from pSVA3919, pSVA5031, and pSVA5611, respectively, for cloning into pTA962-CetZ containing vectors and pHVID132. |
| XbaI_ArlD1 | GGCGGCTCTAGAATGTACCTGGACCCG | Forward primer used to amplify the ArlD1-GFP ORF, introducing an XbaI site to the 5' end for cloning into pHVID132. |
| XbaI_GFP | GGCGGCTCTAGAATGAGTAAAGGAGAAG | Forward primer used to amplify the GFP-CheW1 ORF, introducing an XbaI site to the 5' end for cloning into pHVID132. |
| XbaI_CheY | GGCGGCTCTAGAATGGCAAGCAAGGT | Forward primer used to amplify the CheY-GFP ORF, introducing an XbaI site to the 5' end for cloning into pHVID132. |

**References**


1. Legerme G, Yang E, Esquivel RN, Kiljunen S, Savilahti H, Pohlschroder M: **Screening of a Haloferax volcanii transposon library reveals novel motility and adhesion mutants**. *Life* 2016, **6**(4):41.
2. Duggin IG, Aylett CH, Walsh JC, Michie KA, Wang Q, Turnbull L, Dawson EM, Harry EJ, Whitchurch CB, Amos LA: **CetZ tubulin-like proteins control archaeal cell shape**. *Nature* 2015, **519**(7543):362.
3. Allers T, Ngo H-P, Mevarech M, Lloyd RG: **Development of additional selectable markers for the halophilic archaeon Haloferax volcanii based on the leuB and trpA genes**. *Applied and environmental microbiology* 2004, **70**(2):943-953.
4. Li Z, Kinosita Y, Rodriguez-Franco M, Nußbaum P, Braun F, Delpech F, Quax TE, Albers S-V: **Positioning of the motility machinery in halophilic archaea**. *MBio* 2019, **10**(3):e00377-00319.
5. Quax TE, Altegoer F, Rossi F, Li Z, Rodriguez-Franco M, Kraus F, Bange G, Albers S-V: **Structure and function of the archaeal response regulator CheY**. *Proceedings of the National Academy of Sciences* 2018, **115**(6):E1259-E1268.
6. Ithurbide S, de Silva RT, Brown HJ, Shinde V, Duggin IG: **A vector system for single and tandem expression of cloned genes and multi-colour fluorescent tagging in Haloferax volcanii**. *Microbiology* 2024, **170**(5):001461.
7. Allers T, Barak S, Liddell S, Wardell K, Mevarech M: **Improved strains and plasmid vectors for conditional overexpression of His-tagged proteins in Haloferax volcanii**. *Applied and environmental microbiology* 2010, **76**(6):1759-1769.
